# Dependence of cell fate potential and cadherin switching on primitive streak coordinate during differentiation of human pluripotent stem cells

**DOI:** 10.1101/2025.01.31.635963

**Authors:** Ye Zhu, Aryeh Warmflash

## Abstract

During gastrulation, the primitive streak (PS) forms and begins to differentiate into mesendodermal subtypes. This process involves an epithelial-mesenchymal transition (EMT), which is marked by cadherin switching, where E-Cadherin is downregulated, and N-Cadherin is upregulated. To understand the relationships between differentiation, EMT, and cadherin switching, we made measurements of these processes during differentiation of human pluripotent stem cells (hPSCs) to PS and subsequently to mesendoderm subtypes using established protocols, as well as variants in which signaling through key pathways including Activin, BMP, and Wnt were modulated. We found that perturbing signaling so that cells acquired identities ranging from anterior to posterior PS had little impact on the subsequent differentiation potential of cells but strongly impacted the degree of cadherin switching. The degree of E-Cadherin downregulation and N-Cadherin upregulation were uncorrelated and had different dependence on signaling. The exception to the broad potential of cells throughout the PS was the loss of definitive endoderm potential in cells with mid to posterior PS identities. Thus, cells induced to different PS coordinates had similar potential within the mesoderm but differed in cadherin switching. Consistently, E-Cadherin knockout did not alter cell fates outcomes during differentiation. Overall, cadherin switching and EMT are modulated independently of cell fate commitment in mesendodermal differentiation.

## Introduction

During mammalian gastrulation, the primitive streak (PS) forms and subsequently differentiates into endoderm and mesoderm subtypes. This process involves an epithelial-mesenchymal transition (EMT), during which PS cells change from epithelial to mesenchymal morphology and begin to migrate. EMT is marked by cadherin switching, where E-Cadherin (E-Cad; also known as Cadherin 1 or CDH1) is downregulated, and N-Cadherin (N-Cad; also called Cadherin 2 or CDH2) is upregulated (reviewed by Amack, 2021). In addition to cell adhesion, cadherins were also shown to play a crucial role in fate commitment during neural differentiation. E-Cad downregulation actively promotes neural differentiation (Malaguti et al., 2013), while N-Cad expression helps stabilize neural fate in the neural epithelium (Punovuori et al., 2019). Nonetheless, the role of cadherin switching in mesendodermal fate commitment remains unclear.

Protocols to induce human pluripotent stem cells (hPSCs) to PS and mesendodermal subtypes, such as definitive endoderm (DE), paraxial mesoderm (PM), and lateral mesoderm (LM), have been well established using combinations of Activin/Nodal/TGFβ (hereafter Activin), BMP, Wnt, and FGF/ERK signaling activation and inhibition (Burridge et al., 2014; Gertow et al., 2013; Cheung et al., 2012; Umeda et al., 2012; Patsch et al., 2015; Bernardo et al., 2011; Martyn et al., 2019). Differentiation to any of these subtypes can be achieved with two days of differentiation from hPSCs, with the first day corresponding to PS induction and the second corresponding to further specialization to PM, LM, or DE. Protocols to induce PS include FGF and Wnt activation along with either Activin (anterior PS), BMP (posterior PS), or both (mid PS). Protocols for DE induction on day 1 are similar to those for APS induction with increased Activin and decreased Wnt stimulation (Loh et al., 2014). Cells are thought to be partially committed to mesodermal subtypes following PS differentiation (Hsu et al., 2018; Loh et al., 2014; Loh et al., 2016; Loh et al., 2016; Nakanishi et al., 2009). DE differentiation on day 2 involves continued Activin stimulation with BMP inhibition while the combination of Wnt activation and BMP inhibition induces PM from APS. Conversely, BMP activation combined with Wnt inhibition induces LM from MPS (Loh et al., 2016) (Fig. 1A).

**Figure 1.**
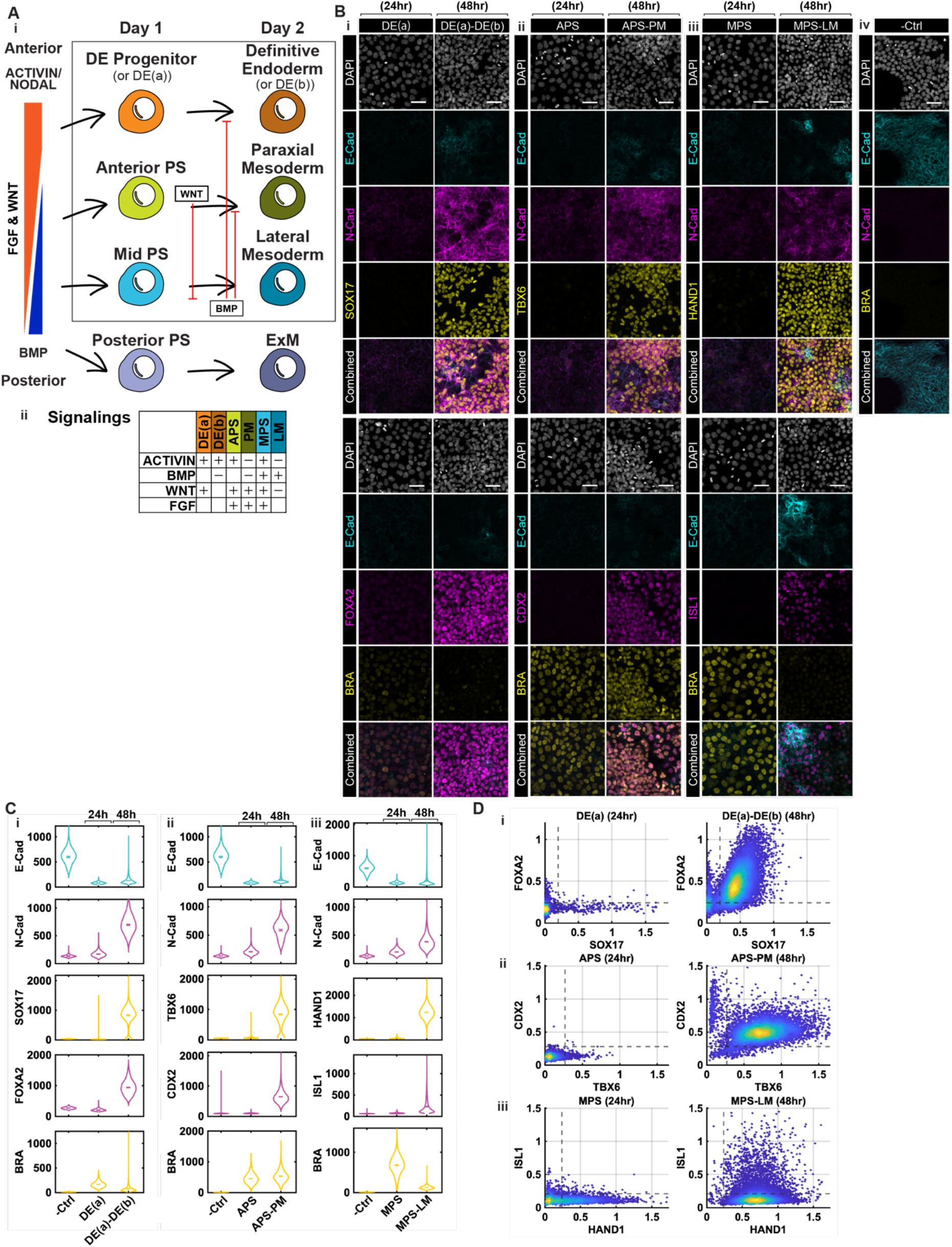
Fate marker and cadherin expression during established protocols for DE, PM, and LM. **A**. Schematic of established protocols for mesendoderm induction from hPSCs through the Activin, BMP, Wnt and FGF signaling pathways. **B**. Example confocal immunofluorescent images of hPSCs following the indicated treatments. Fate-specific markers are used: BRA for PS; SOX17 and FOXA2 for DE(a) and (b); TBX6, and CDX2 for APS and PM; HAND1, and ISL1 for MPS and LM, and the cadherin markers E-Cad, and N-Cad. Scale Bars: 50 µm. **C**. Quantification of fluorescent intensity of the indicated markers following the indicated protocols. **D**. Co-expression scatter plots of normalized mean fluorescence intensity in each cell (relative to DAPI) showing FOXA2 vs. SOX17 for DE(a), and (b); CDX2 vs. TBX6 for APS and PM; ISL1 vs. HAND1 for MPS and LM. (N = 6 images per condition.)

While these protocols have been extensively used, the degree to which the potential of different PS populations differs as well as the relationship between signaling and cell fate, on the one hand, and EMT and cadherin switching on the other, remain unclear. To address this gap, we investigated the potency of different PS populations and cadherin switching dynamics in differentiating hPSCs. We found that different regions of the PS maintain similar potency within the mesoderm, indicating that spatial coordinates within the PS do not rigidly dictate mesodermal fate commitment. While anterior PS cells retained potency for both endoderm and mesoderm, mid to posterior PS lost differentiation potential toward definitive endoderm while retaining broad potency within the mesoderm. The degree of cadherin switching did correlate with PS position and was regulated by signaling independently of fate commitment. Further we observed that E-Cad downregulation and N-Cad upregulation were not strongly correlated and occurred independently during mesendoderm differentiation. Finally, loss of epithelial and gain of mesenchymal properties and cadherin switching were also more independent from each other than previously appreciated so that cells could acquire mesenchymal properties even without cadherin switching. Consistently, knocking out E-Cad resulted in only minor effects on mesendodermal differentiation. Taken together, our results suggest that the processes of mesendoderm cell differentiation, EMT, and cadherin switching are largely decoupled and that position within the PS determines the degree of cadherin switching more than cell fate potency.

## Results

### Generation of mesendodermal cell types with established protocols

Using previously reported protocols (Fig. 1A), we differentiated hPSCs into PS subtypes and mesendodermal lineages, generating definitive endoderm (DE), paraxial mesoderm (PM), and lateral mesoderm (LM) within 48 hours (Fig. 1B, C). In each case, we obtained a largely homogenous population with greater than 80% of the cells expressing expected markers: SOX17^+^and FOXA2^+^for DE, TBX6^+^and CDX2^+^for PM, and HAND1^+^for LM. In the case of LM, we observed heterogeneity in ISL1 expression with 20% of cells expressing this marker (Fig. 1D).

To understand the heterogeneity within the LM population, we stained for additional markers associated with extraembryonic (GATA3) and cardiac (NKX2.5) fates (Fig. S3A-D). We found that nearly all ISL1^high^ cells are GATA3^+^, compared to only a minority of HAND1^+^cells expressing GATA3. Notably, the ISL1^high^ cells are also E-Cad positive. Thus, the presence of a small extraembryonic population (HAND1^+^/IS1^high^/GATA3^+^/E-Cad^+^) was identified, in addition to the main LM population (HAND1^+^/ISL1^Low^/E-Cad^−^). This population may be either amnion or extraembryonic mesoderm, both of which express GATA3 (Chen et al., 2024; Chhabra and Warmflash, 2021). On the other hand, NKX2.5 expression did not correlate with high ISL1 and the HAND1 expression in LM: only small percentages of LM cells expressed NKX2.5, however, the LM cells were competent to generate cardiac mesoderm as greater than 50% of cells expressed NKX2.5 when a cardiac differentiation was applied on the third day following LM induction (Fig. S3C, D).

In summary, the DE and PM populations obtained using established protocols were relatively homogeneous, with high co-expression of the key fate markers, while cells subjected to the LM protocol exhibited some heterogeneity and included a small subpopulation of amnion-like cells.

### DE, PM, and LM protocols have different cadherin switching dynamics

We next examined the dynamics of cadherin switching and EMT marker expression during mesendoderm differentiation. E-Cad downregulation was observed at 12 hours after induction across all PS subtypes, marking the initiation of cadherin switching, while N-Cad upregulation was not detected at 12 hours but was evident at 24 hours (Fig. S1A). At 12h, Snail1 expression remained undetectable, suggesting that E-Cad repression can be initiated while Snail1 expression remains low (Fig. S1A). Thus, E-Cad downregulation is an early and consistent event in mesendodermal differentiation, prior to N-Cad or EMT marker activation.

The timing and level of N-Cad upregulation and EMT marker expression varied among the three lineages within the first day of induction (Fig. S1A). DE had the lowest Snail expression, consistent with observations in mouse embryos in vivo that Snail1 expression is limited to the PS and the nascent mesoderm but not DE (Carver et al., 2001; Smith et al., 1992) and that Snail1 is not required for endoderm formation (Scheibner et al., 2021).

By 48 hours, greater than 85% of cells expressed N-Cad in all mesendodermal lineages (Fig. S2Ai, iii). Concurrently, a small subset of approximately 10-20% of cells began re-expressing E-Cad in each lineage with the highest percentage in LM (Fig. S2Ai, iv). Nearly all E-Cad positive cells also expressed N-Cad. Notably, E-Cad^+^extraembryonic cells found in the LM protocol also had high N-Cad expression (Fig. S2Ai). In vivo, the amnion does not express N-Cad, and the extraembryonic mesoderm does not express E-Cad, but our results here indicate cells adopting one of these fates are capable of expressing both cadherins. In short, all populations could be separated by cadherin expression into a majority of cells which were E-Cad−/N-Cad^+^, and a small subgroup of E-Cad^+^/N-Cad^+^cells.

We utilized the markers ZO-1 and VIMENTIN (VIM) to analyze whether cells adopted an epithelial or mesenchymal phenotype (Fig. S2C, D). ZO-1 is localized to tight junctions on the apical side of epithelial cells, including pluripotent cells, while VIM is an intermediate filament protein expressed in mesenchymal cells (Yang et al., 2020). At 24 hours of differentiation, PS cells lost epithelial characteristics with approximately 10% of cells expressing ZO-1 compared to 90% in the pluripotent state. VIM expression remained low, however, with levels comparable to the pluripotent controls (Fig. S2B). By 48hrs, VIM was elevated in all mesendodermal lineages with cells co-expressing N-Cad. Snail dynamics differed between the lineages with downregulation in LM and continued expression in PM (Fig. S1A, C). N-Cad positive cells lost expression of E-Cad at the junctions and appeared mesenchymal in nature.

These observations suggest that while DE, PM, and LM share a similar initiation of EMT and cadherin switching, the subsequent dynamics differ among the lineages, particularly concerning Snail expression dynamics and the fraction of cells which re-express E-Cad.

### Cells at Different Coordinates in the Primitive Streak Have Similar Potency in Mesoderm

We then investigated whether cells differentiation to different positions within the PS, as represented by the treatments APS, MPS, and PPS, exhibit distinct potencies toward PM and LM. As the PS protocols differ in the inclusion of Activin and BMP, we also included a control group without addition of either ligand, labeled as the -TGFβ group (Fig. 2A).

**Figure 2.**
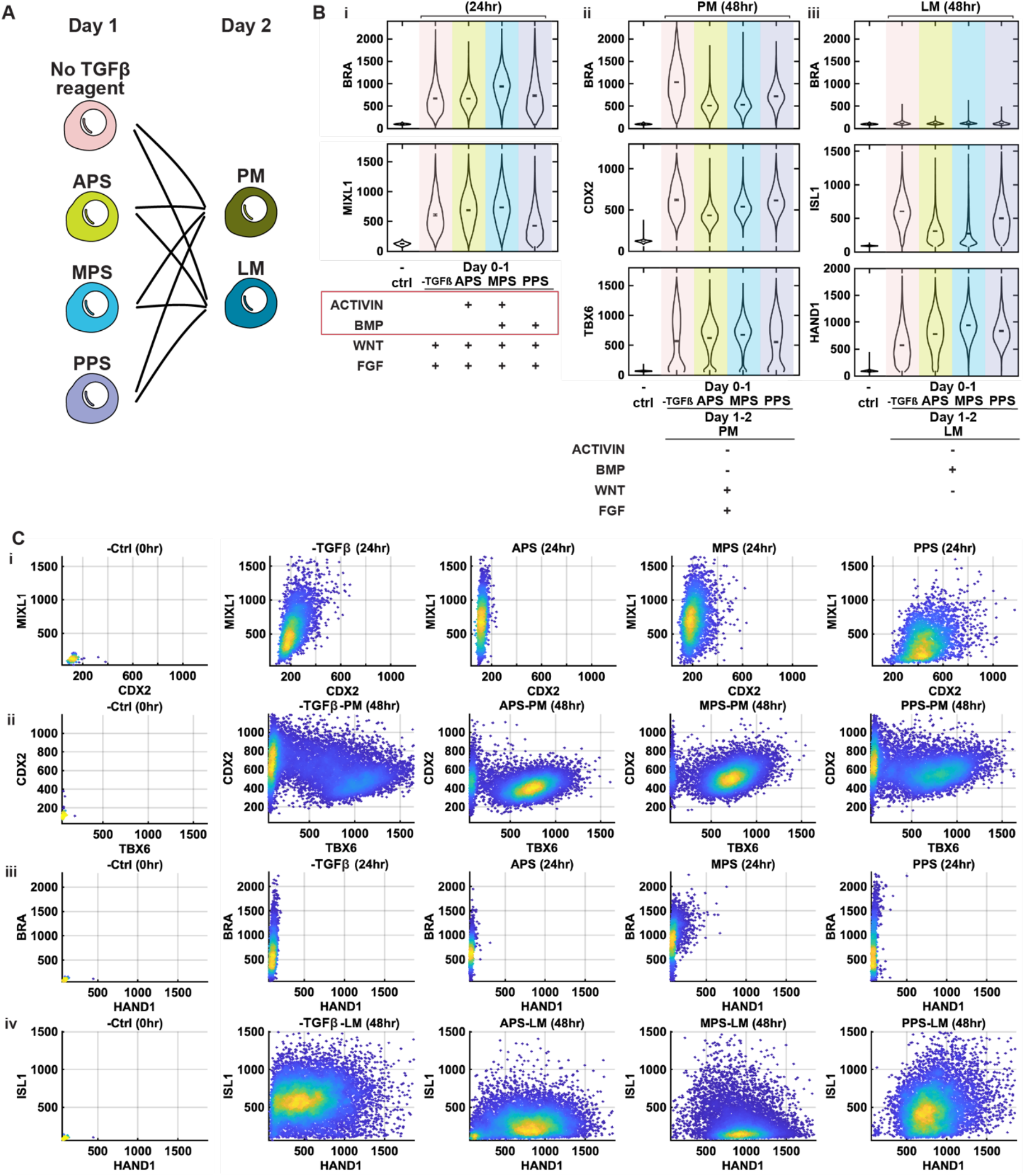
Cells at Different Coordinates in the PS Have Similar Mesodermal Potency. **A**. Schematic of the experimental design testing PM and LM potency of APS, MPS, PPS, and the treatment with no Activin A or Bmp4 (denoted the -TGFβ group). **B**. Quantification of markers of primitive streak (BRA, and MIXL1), PM(BRA, CDX2, and TBX6) and LM (BRA, ISL1, HAND1) from confocal microscopic images of cells with the indicated treatments and fixed at the time indicated above the plots. **C**. Co-expression scatter plots of fate markers for the indicated treatments and fixation times.

All four groups activated BRA and MIXL1 by 24 hours (Fig. 2Bi). Upon further induction toward PM, all groups expressed TBX6 and CDX2, and when directed toward LM, HAND1 expression was robust, indicating that APS, MPS, PPS, and -TGFβ groups all retained potency for PM and LM fates (Fig. 2Bii, iii), although quantitatively there were minor differences in the distribution of marker expression depending on the PS induction protocol (Fig. 2C). Both the -TGFβ and PPS groups were deprived of Activin A during PS induction, and these groups displayed lower TBX6 and higher CDX2 expression in PM induction at 48 hours, as well as higher ISL1 expression in LM induction. These findings suggest that Activin signaling during the first 24 hours helps prevent cells from committing to extraembryonic fates while enhancing potential for mesodermal fates (Fig. 2B), consistent with data from modulated Activin signaling during micropatterned differentiation (Jo et al., 2022). Taken together, our data suggest that once cells adopt primitive streak fates, they have similar potential for different fates within the mesoderm regardless of the coordinate, however, signaling that varies with position in the PS may influence the decision between extraembryonic and PS fates with extraembryonic fates more suppressed at anterior coordinates.

### Endoderm Progenitors in Primitive Streak Exhibit Differential Potency Towards Mesodermal and Endodermal Fates

We assessed the endodermal potency of different PS subtypes and included a treatment which is similar to APS but has been optimized for endoderm differentiation (Loh et al., 2014) (marked as DE(a), the second day is marked as DE(b)). Cells treated with DE(a) activated BRA, the PS/mesodermal progenitor marker, albeit at the lowest level compared to the other PS subtypes (Fig. 3B). At this stage, expression of the endodermal marker SOX17 remained undetectable. Upon extending the treatment of different PS subtypes with DE(b) in the second day, the DE(a)-DE(b) and APS-DE(b) groups showed similar co-expression of the endodermal markers SOX17 and FOXA2 (Fig. S4ii). In contrast, MPS and PPS largely failed to express SOX17 and FOXA2 (Fig. 3B). These findings suggest that the DE(a) and APS subtypes retain potency for both DE and PM, whereas the mid-to-posterior PS subtypes lose their potency for DE.

**Figure 3.**
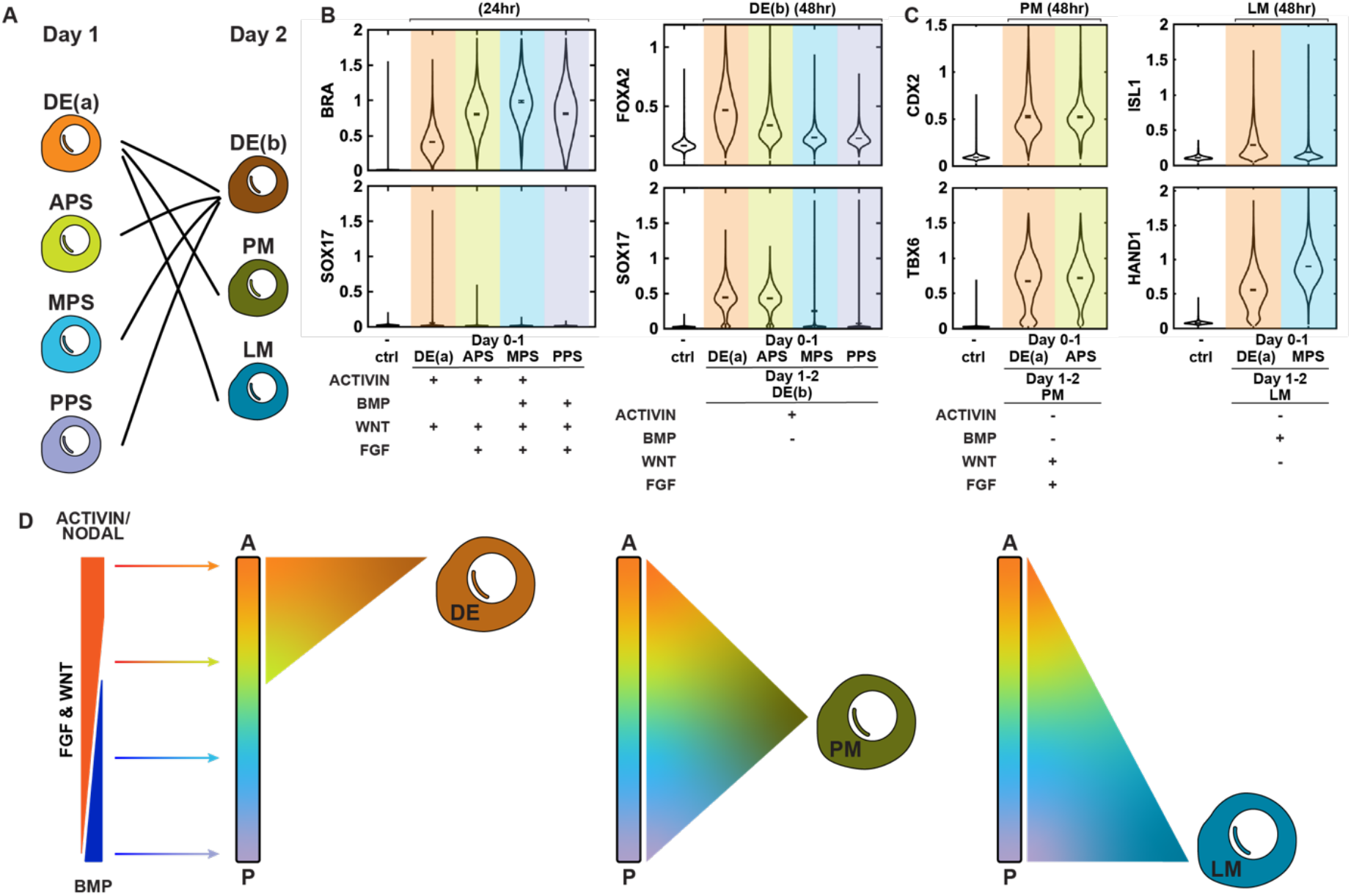
Anterior PS cells have broad potential while mid and posterior PS cells lose DE potential. **A**. Schematic of the experimental design testing the DE potency of APS, MPS, PPS, and mesodermal potency of DE(a) treated cells. **B**. Quantification of endodermal markers from confocal microscopic images at the following the indicated treatments and fixation times. **C**. Quantification of PM and LM comparing standard protocols and those in which an endodermal differentiation protocol is followed on day 1. **D**. Schematic indicating the range of PS cell types with the potential to give rise to each further differentiated cell type.

We tested the mesodermal potential of cells treated with the endoderm optimized DE(a) protocol. By extending the treatment with PM on day 2, DE(a)-PM exhibited an identical TBX6/CDX2 expression distribution as the original APS-PM treatment. Similarly, DE(a)-LM also activated HAND1 expression, confirming the broad mesodermal potency of DE(a) cells (Fig. 3C).

In summary, cells from different coordinates in the PS (including endodermal progenitors) presented similar potency toward mesodermal subtypes. However, cells at the mid-to-posterior end of the PS lost potency toward endoderm (Fig. 3D).

### The Coordinate Within the Primitive Streak Strongly and Separately Affects E-Cad Downregulation and N-Cad Upregulation

While mesendodermal treatment groups exhibited similar potency and fate commitment across lineages, the dynamics of cadherin switching and EMT progression varied significantly across the PS subtypes from anterior to posterior. To explore these differences, we examined expression of Snail, E-Cad, and N-Cad in the APS, MPS, PPS, and -TGFβ groups directed towards PM and LM fates.

During the initial 24 hours as cells exited pluripotency, Activin downregulated while BMP upregulated E-Cad so that the lowest levels of E-Cad were found in APS and the highest levels in PPS. In contrast, both Snail and N-Cad expression were highest in the condition with both Activin and BMP treatment, suggesting that both signals contribute to upregulating these factors (Fig. 4A, S6A). At 48 hours, these trends largely persisted with E-Cad highest in the posterior conditions while almost completely lost in APS and MPS conditions, and N-Cad and Snail highest in the mid to posterior regions (Fig. S6B, C). Both N-Cad and Snail expression were higher in PM than LM across all PS coordinates, which may reflect a switch in the role of BMP from activating to repressing these factors between day 1 and day 2. This will be explored in more detail below.

**Figure 4.**
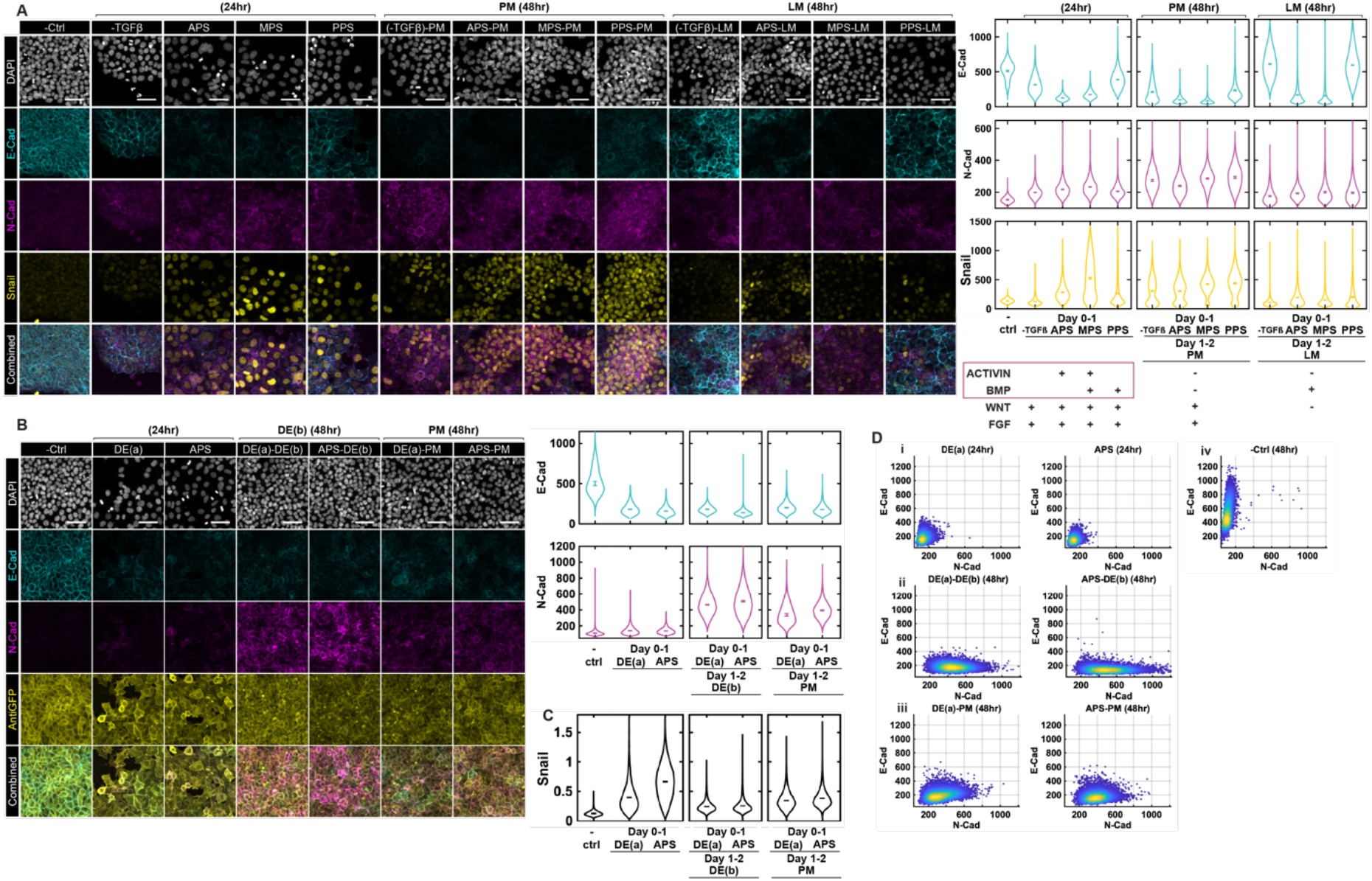
Cadherin switching in mesendodermal differentiation is independent of fate commitment. **A**. Example confocal immunofluorescent images and quantification based on fluorescent intensity of E-Cad, N-Cad, and Snail for the indicated treatments and fixation times. (**B, C)**. Example confocal immunofluorescent images and quantification based on fluorescent intensity of E-Cad and N-Cad (B) or Snail (C) comparing standard protocols for PM with those in which DE(a) was followed on day 2.) **D**. Co-expression scatter plots of E-Cad and N-Cad comparing using DE(a) or APS on day 1 of differentiation followed by either PM or DE(b) (N = 6 images per condition.)

In absence of all added TGFβ ligands, both N-Cad and Snail activation were impaired. In the -TGFβ-LM condition, these factors were completely absent while E-Cad was strongly expressed, as BMP upregulated E-Cad expression but not Snail or N-Cad expression on day 2 (Fig. 4A, S6C). In PM, Snail and N-Cad were upregulated while E-Cad was downregulated (Fig. S6B), likely in response to the strong Wnt signal activation on day 2 in this condition (see below).

We also compared DE(a) and APS followed by either DE(b) or PM induction on day 2. In general, cells with APS yielded lower E-Cad and higher Snail and N-Cad compared to DE(a) (Fig. 4B-D). This likely reflects the higher concentration of CHIR in APS compared to DE(a) as Wnt signaling promotes both cadherin switching and Snail expression (see below).

Taken together these data suggest that signals which vary with the PS coordinate affect cadherin switching and Snail expression on both the first and second day. On day 1, during PS induction, Activin downregulates, while BMP maintains, E-Cad expression, while both of these signals upregulate N-Cad and Snail expression. Thus, while N-Cad and Snail expression closely mirror each other, the expression of E-Cad and N-Cad are largely decoupled due to their differential regulation by Activin signaling. By day 2, the expression of all three of these markers can vary within cells of the same cell fate, as their expression, but not the cell fate, depends on the day 1 conditions.

### Transcriptome analysis confirms broad potential of PS progenitors

To validate our results on PS potential at the transcriptome level, we performed RNA sequencing of samples differentiated to the four primitive streak populations (DE(a), APS, MPS, and PPS) at 24 hours as well as each of these populations further differentiated to LM, PM, or DE(b). Hierarchical clustering of all samples grouped samples based on the day 2 treatment without regard to the PS induction protocol on day 1, indicating that generally all the PS progenitors are capable of yielding populations with similar transcriptomes when further differentiated (Fig. 5A). Consistently, performing PCA and coloring samples on the PC1 vs PC2 plot by day 2 treatment showed coherent groups of conditions, while coloring based on day 1 treatment did not (Fig. 5B).

**Figure 5.**
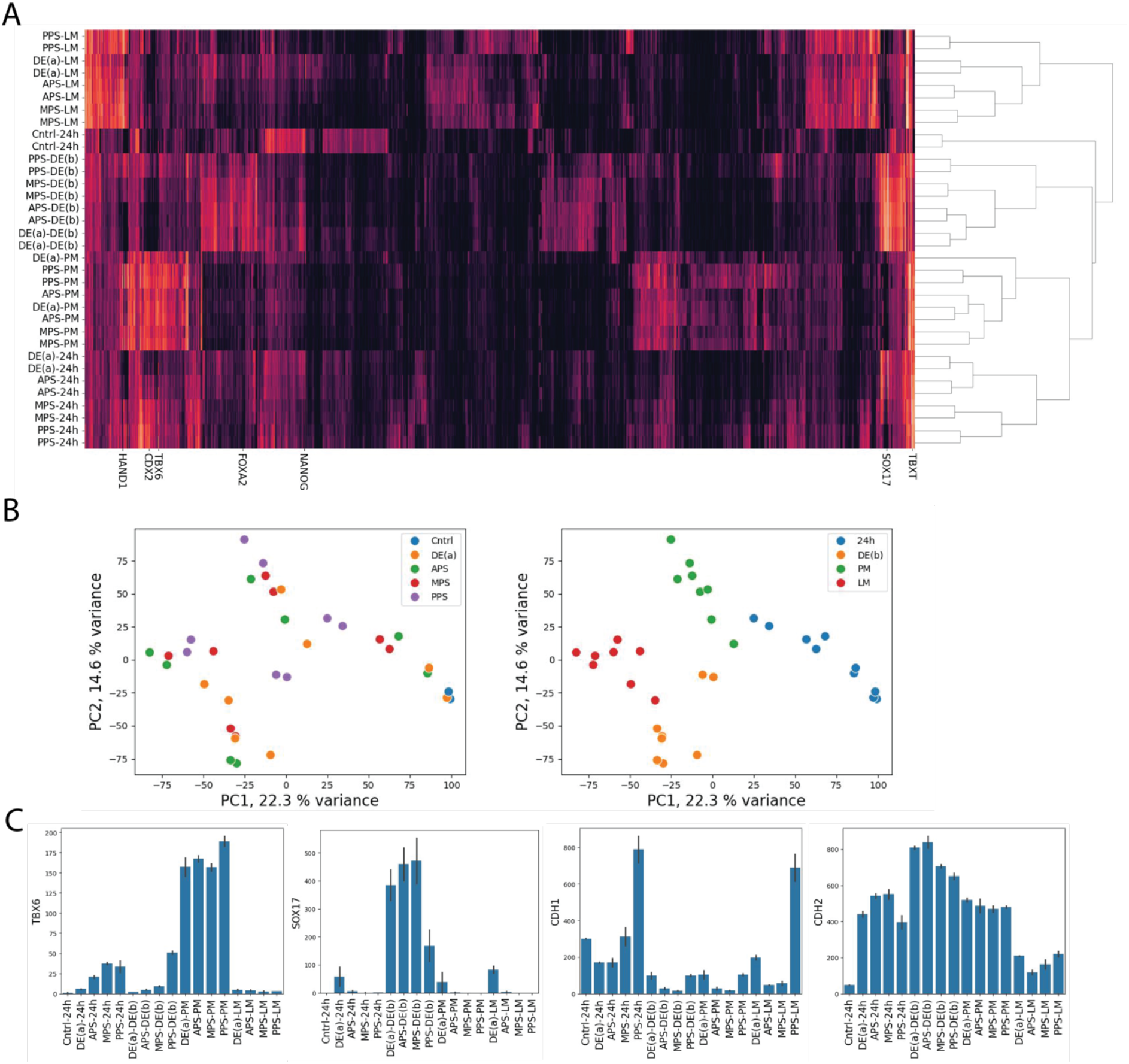
Transcriptomic analysis confirms broad potential of PS cells. **A**. Hierachical clustering of RNA sequencing data. Some of the marker genes used in this study are indicated. **B**. Principal component plot of RNA sequencing data colored by either the PS induction protocol performed on day 1 (left) or the differentiation protocol on day 2 (right). C. Expression of individual genes in RNA sequencing samples.

At the level of individual genes, we confirmed that fate markers such as TBX6 were expressed at similar levels regardless of the initial PS differentiation (Fig. 5C). SOX17 was also expressed at similar levels in all PS protocol followed by DE(b) except for PPS-DE(b), consistent with our results above that posterior PS loses the potential for DE differentiation. The cadherin genes CDH1 and CDH2 showed variability within each fate with in general highly levels of CDH1 (E-Cad) expression when the initial treatment was either DE(a) or PPS compared with the APS or MPS treatments. As in the immunostaining data, CDH2 (N-Cad) showed different trends depending on cell fates with levels increasing with posterior PS in LM differentiation but decreasing in DE or PM treatment (Fig. 5C).

### Activin Promotes, and BMP Inhibits, Cadherin Switching During MPS-LM Induction

To better visualize and quantify cadherin dynamics in time during mesendodermal differentiation, we generated a dual-reporter hPSC line, with fluorescent tags at the endogenous loci for E-Cad and N-Cad and a DOX-inducible membrane fluorescent marker for cell identification (ESI-017 E-Cad:mCitrine N-Cad:mCherry CAAX:mCerulean, Fig. 6A, Movie 1). This reporter cell line maintained pluripotency and exhibited mesendodermal induction potency comparable to the wild-type (WT) parental line (Fig. S7B-D). We focused on modulating the MPS-LM protocol as MPS contains all four signals involved in the primitive streak.

**Figure 6.**
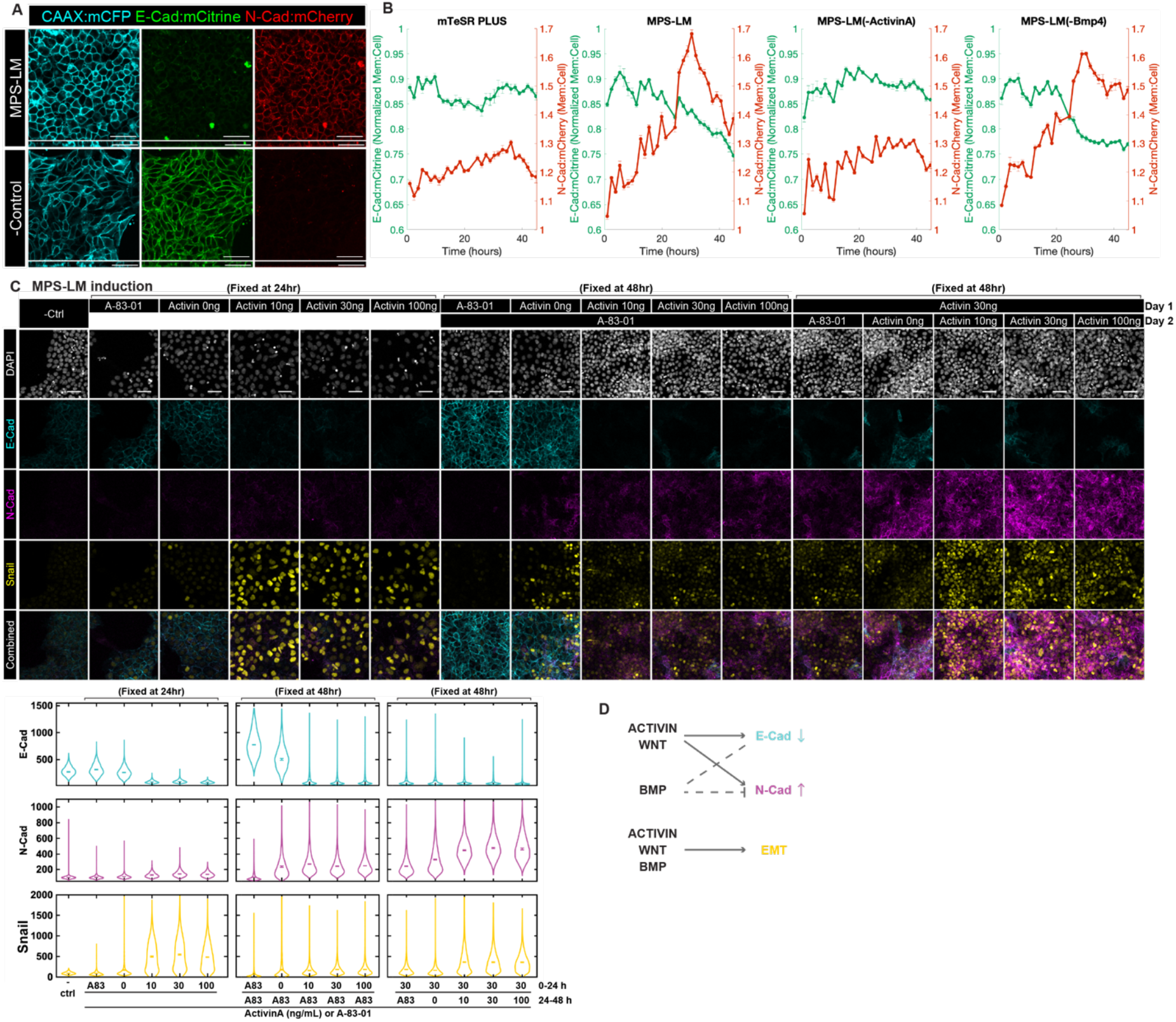
Cadherin switching is regulated by Activin and BMP signaling. **A**. Example confocal microscopy images of live ESI017-CDH1:mCitrine-CDH2:mCherry-CAAX:mCerulean reporter cell line after 48hr MPS-LM treatment or mTeSR Plus control treatment. Scale bar: 50 µm. **B**. Quantification from 0-44hr of the fluorescent intensity of CDH1:mCitrine and CDH2:mCherry for cell with the indicated treatments 0-44 hr. (N = 3 positions per condition.) **C**. Example confocal immunofluorescent images and quantification based on fluorescent intensity of E-Cad, N-Cad, and Snail for cells that underwent MPS or MPS-LM induction with either A-83-01 (1 µM) or various Activin A doses (0, 10, 30, and 100 ng/mL) and fixed by 24hr (N= 7 images per condition.) **D**. Schematic of the relationship between ACTIVIN, BMP and WNT signaling with E-Cad downregulation, N-Cad upregulation, and EMT activity

Live-cell imaging during MPS-LM induction captured the progression of cadherin switching. E-Cad downregulation initiated early, while the reporter allowed us to capture gradual upregulation of N-Cad that began almost immediately upon differentiation and continued throughout the first day. N-Cad upregulation accelerated on the second day and peaked at around 30 hours, and then downregulated and stabilized by 48 hours (Fig. 6B). These rapid changes likely reflect the switch from MPS to LM media and highlight the dynamic nature of cadherin expression in response to signaling during mesodermal differentiation.

To elucidate the roles of Activin and BMP signaling in this process, the MPS-LM induction protocol was modified by withdrawing Activin A or Bmp4 (Fig. 6B). Notably, Activin had a stronger effect on cadherin dynamics than Bmp signaling. Removing Activin A impaired both E-Cad downregulation and N-Cad upregulation, demonstrating its critical role in promoting cadherin switching. In contrast, withdrawing Bmp4 had minimal impact on E-Cad downregulation but caused N-Cad to stabilize at a higher level by 48 hours, confirming the inhibitory role of Bmp on N-Cad upregulation on the second day of differentiation, consistent with our results above. In summary, Activin and Bmp have opposing roles in regulating cadherin switching during lateral mesodermal induction, where Activin promotes cadherin switching and Bmp selectively inhibits N-Cad expression on day 2 (Fig. 6D).

### Roles of Activin, BMP, and Wnt doses during differentiation

We further investigated the effects of Activin signaling in initiation of EMT and cadherin switching, as well as fate specification for LM induction by either adding the Activin inhibitor A-83-01 or varying the dose of Activin A (0, 10, 30, or 100 ng/mL; with 30ng/mL Activin A being the original dosage).

On day 1, withdrawing Activin A or adding the inhibitor A-83-01 impaired E-Cad downregulation and Snail and N-Cad upregulation (Fig. 6C), emphasizing the critical role of exogenous Activin signaling in initiating EMT and downregulating E-Cad during early PS induction. By continuing these groups with the original second-day LM induction protocol, we observed that, when Activin was included, varying the dosage did not significantly alter LM fate commitment (HAND1^+^/ISL1^low^) or cadherin dynamics (Fig. 6C, S8). Inhibition of Activin/Nodal signaling caused E-Cad to remain highly expressed and a failure to express N-Cad, while simply removing the exogenous Activin caused co-expression of E-Cad and N-Cad across the population. The effects on cell fate were less pronounced, however, the absence of Activin signaling on the first day promoted GATA3 and ISL1 expression, increasing the fraction of cells in the extraembryonic lineage (Fig. S8A). Nonetheless, there was still substantial mesoderm differentiation as reflected in BRA expression, with the resulting population a mixture of amnion and mesoderm (Fig. S8B). These results show Activin signaling is required for both E-Cad downregulation and N-Cad upregulation while it quantitatively biases cells to mesodermal rather than amnion lineages, but is not strictly required.

Removing A83 from the day 2 LM protocol significantly enhanced N-Cad expression without changing the expression of fate markers. Adding exogenous Activin further enhanced N-Cad expression, and diverted cells away from the LM lineage as reflected in a dose-dependent loss of HAND1 expression (Fig. 6C, S8). In summary, Activin signaling on the first day is essential for initiating EMT, downregulating E-Cad, and suppressing extraembryonic fates, while its role on the second day is primarily to promote N-Cad upregulation. High levels of Activin signaling during day 2 are not consistent with LM fate commitment.

We also investigated modulating BMP signaling levels during the MPS-LM protocol and found only minor effects of BMP dosage on cell fate or cadherin switching. Most significantly, inhibiting BMP on day 1 resulted in lower expression of both N-Cad and E-Cad expression while somewhat reducing potency of cells for LM as reflected in a fraction of cells failing to express HAND1. In contrast, inhibiting BMP on day 2, compromised LM induction but had only minor impacts on cadherin switching (Fig. S9A, B).

We similarly investigated the role of Wnt signaling in the APS-PM protocol. Removing CHIR or adding IWP on day 1 compromised E-Cad downregulation, Snail and N-Cad upregulation, and cell fate specification, while inhibiting it on day 2 impaired PM differentiation with smaller effects on cadherin or Snail expression (Fig. S10A, B). Thus, Wnt signaling is essential for all aspects of PS formation, EMT and cadherin switching on day 1, and for PM differentiation on day 2.

### E-Cadherin Knockout Has Minor Impacts on Fate Induction and Lineage-specific Cadherin Dynamics

Our results above suggest that cell fate acquisition and EMT are largely independent of Cadherin switching while N-Cad upregulation and E-CAD downregulation are also separate processes. To validate these results functionally, we created an E-CAD KO cell line through CRISPR-Cas9 RNP delivery. The loss of E-Cad was validated via sequencing and immunostaining (Fig. 7A, S11A). Notably, E-Cad^−/−^ cells localized ZO-1 expression to the apical side, showing an epithelial morphology with apical-basal polarity (Fig. 7A). The pluripotent state (SOX2^+^, OCT4^+^, and NANOG^+^)of the E-Cad^−/−^ cells remained comparable to that of WT cells (Fig. S11C).

**Figure 7.**
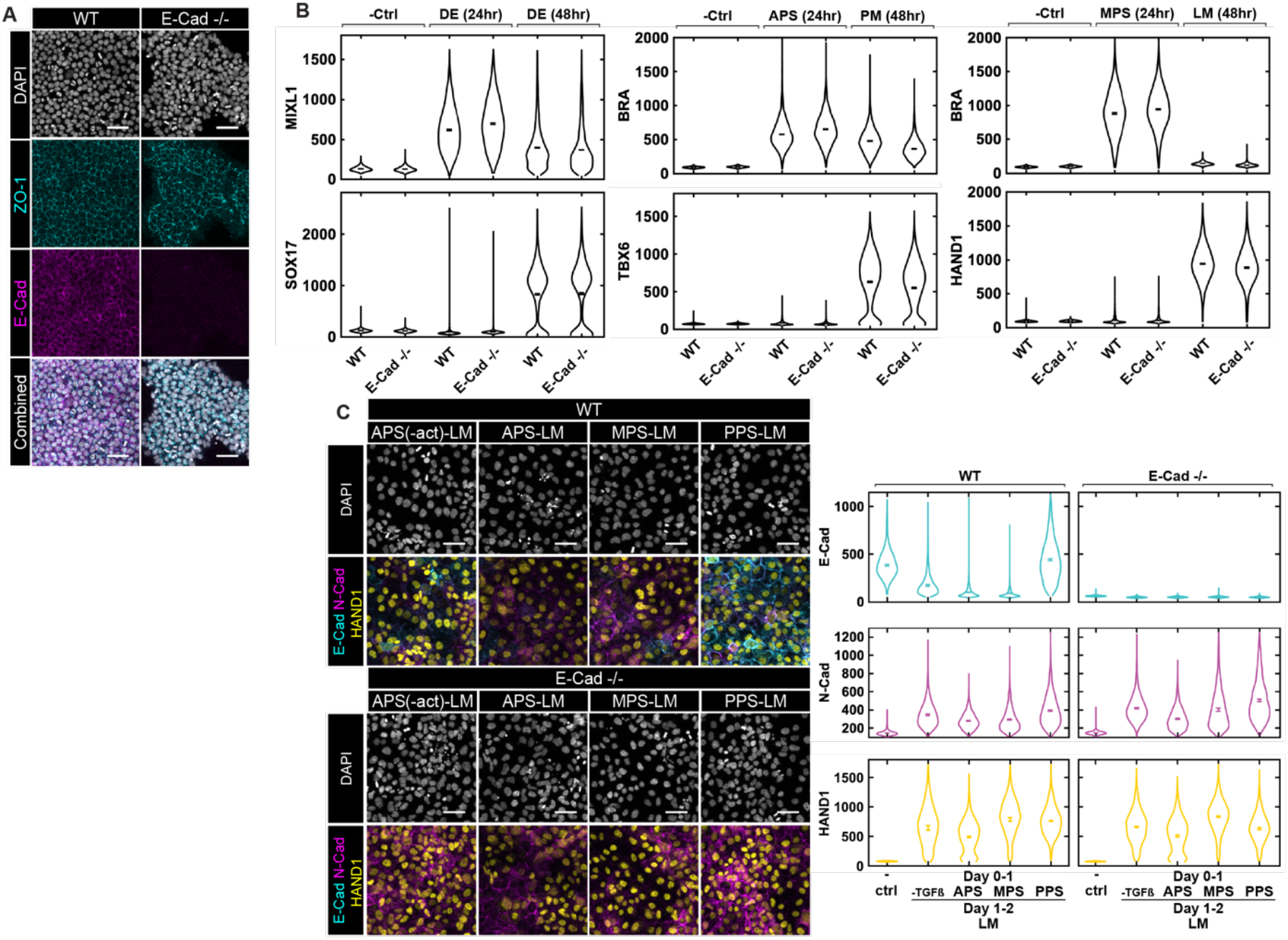
CDH1 (E-Cad) knock out has minor effects on fate commitment in mesendoderm differentiation. **A**. Example confocal microscopic images of ESI017 wild type (WT) and E-Cad ^−/−^ (maintained in mTeSR Plus) immunostained for DAPI, ZO-1 and E-Cad. **B**. Quantification based on fluorescent intensity of ESI017 wild type (WT) and E-Cad ^−/−^ following the indicated standard protocols. – Ctrl: negative control that is maintained in mTeSR Plus and fixed at 0hr. (N = 8 images per condition.) **C**. Example confocal microscopic images and quantification of E-Cad, N-Cad, and fate markers for WT and E-Cad ^−/−^ cells with varying PS induction treatment on day 1 and induced to LM on day 2. Scale Bars: 50 µm. – Ctrl: negative control that is maintained in mTeSR Plus. (N = 5 images per condition.)

Differentiation of E-Cad^−/−^ cells was similar to wildtype in PS induction (MIXL1, BRA), and the subsequent DE (SOX17), PM (TBX6), and LM (HAND1) differentiation (Fig. 7B). E-Cad^−/−^ differentiated to different PS subtypes maintained lineage potency toward both PM and LM fates just as WT cells, with indistinguishable TBX6 and HAND1 expression between the cell lines (Fig. 7C, S13). These results suggest that removal of E-Cad does not affect the cell fate specification processes directly.

We also examined N-Cad and Snail1 expression in E-Cad KO cells and noted minor differences with wildtype (Fig. S12). Snail expression was slightly but consistently lower in E-Cad KO compared to wildtype cells, while variable differences were noted in N-Cad expression. In particular, E-Cad KO cells showed reduced N-Cad during DE differentiation but increased N-Cad during LM differentiation. Taken together these results suggest that E-Cad is not critical for PS and subsequently endodermal and mesodermal differentiation. It is also not required for Snail or N-Cad expression, but its loss changes the quantitative expression of these factors in a manner that depends upon the lineage.

## Discussion

In this study, we used hPSCs to examine the dynamics of differentiation, cadherin switching, and EMT during mesendoderm differentiation. The first day of the protocol involves exit from pluripotency and upregulation of PS markers and is generally considered to specify the coordinate within the primitive streak. The second day’s treatment typically dictates the subset of mesendoderm such as endoderm, lateral mesoderm or paraxial mesoderm. We found that cells with different PS coordinates had broad and overlapping potential so that, with a few exceptions, the cell fate was determined by the treatment on the second day. In contrast, cadherin expression depended on signaling conditions on both days of treatment and was more graded with cells adopting each cell fate capable of expressing varying levels of both E-Cad and N-Cad. Moreover, the degree of E-Cad downregulation and N-Cad upregulation could be varied independently, and cadherin expression was not strictly linked to expression of EMT regulators such as Snail or to markers of epithelial or mesenchymal properties such as ZO-1 and VIM. Taken together, our results suggest the events that occur in a coordinated fashion during gastrulation in vivo are regulated separately and can be modulated independently in vitro.

In vivo fate mapping has shown that mesendoderm subtypes emerge from particular PS coordinates, such as lateral mesoderm from the posterior PS and endoderm from the anterior PS. Based on these known lineages, in vitro protocols were developed to regionalize cells within the PS on the first day before subsequent differentiation to particular mesendodermal fates on later days (Loh et al., 2016), however, this study did not compare different PS coordinates for their potency to generate downstream subtypes. Another study compared mesoderm induced with Activin and BMP to that induced by Wnt and concluded that these have different potential (Mendjan et al., 2014), but neither of these conditions correspond directly to different primitive streak coordinates: in the Activin + BMP case, because Wnt was not added, and in the Wnt only case because no TGFβ signaling was included. Moreover, this study concluded that Wnt signaling could be inhibited in the Activin + BMP context without compromising mesoderm induction, whereas our results here (Fig. S10) as well as those from a number of in vivo and in vitro studies (Chhabra et al., 2019; Huelsken et al., 2000; Liu et al., 1999; Martyn et al., 2018) argue that Wnt is indispensable for mesendoderm differentiation. Thus, it is difficult to draw conclusions from this study regarding the potential of cells from different PS coordinates.

Our findings here suggest that cells throughout the primitive steak maintain broad but not identical potential. All PS subtypes we tested were capable of efficiently giving rise to both lateral and paraxial mesoderm. On the other hand, mid to posterior PS cells did not give rise to endoderm. Although not one of our PS conditions, in our studies of BMP dosage, we also found that cells with inhibition of BMP type I receptor kinases with LDN on day 1 were impaired in giving rise to lateral mesoderm. It remains to be determined whether this situation exists in vivo or is an artifact of these culture conditions. Altogether our studies suggest that cells throughout the PS have broad, overlapping, but not universal potential for mesendodermal fates. Our results are more consistent with the idea that AP coordinate and mesendoderm potency exist on a spectrum rather than a rigid delineation between anterior and posterior progenitors that is the assumption of earlier work (Loh et al., 2016; Mendjan et al., 2014).

During gastrulation, mesoderm differentiation in the primitive streak is accompanied by a switch from E-to N-Cadherin expression and acquisition of mesenchymal migratory properties. In Drosophila, embryos lacking either E-CAD, N-CAD, or both undergo normal mesoderm differentiation, patterning and morphogenesis indicating cadherins are not required for either process in this context, however, overexpression of E-Cad can lead to defects due to inhibition of Wnt signaling through sequestration of β-catenin (Schäfer et al., 2014). In mice, evidence from knockout phenotypes suggests that cadherin migration and differentiation may be separable. For example, mesoderm differentiation occurs in EOMES knockout mice but cells fail to undergo EMT, although Snail1 expression is not affected (Arnold et al., 2008). Similarly, mutants for FGFR1 can specify mesoderm but show an accumulation of cells in the primitive streak that indicates a failure of cells to undergo EMT. It was suggested that a pathway in which FGF signaling upregulates Snail1 which in turn downregulates E-Cadherin is important for EMT in this context (Ciruna and Rossant, 2001; Deng et al., 1994; Yamaguchi et al., 1994). In all of these cases, however, mesoderm patterning was also affected with particular subsets of mesoderm enhanced at the expense of others, which may have resulted from dysregulation of signaling. For example, Fgfr1 mutants have attenuated Wnt signaling which can be relieved through blocking of E-Cad (Ciruna and Rossant, 2001). In vitro studies provide an opportunity to decouple the effects of cadherins from changes in the signaling environment.

Our results from in vitro experiments suggest that cadherin switching can be modulated independently of cell fate. For example, cells differentiated to different primitive streak populations (APS, MPS, or PPS) and then subjected to a paraxial mesoderm differentiation protocol on day 2 all display a large fraction of TBX6, CDX2 double positive paraxial mesoderm progenitors (Fig. 3). However, compared with the published APS-PM protocol, the MPS-PM protocol yields stronger N-Cad expression, while the PPS-PM protocol yields high levels of both E-Cad and N-Cad (Fig. 4). These results also suggest that Cadherin switching should not be considered a single coordinated process but rather two separate events, E-Cad downregulation and N-Cad upregulation, which need not occur together. Treatments such as PPS-PM yield high levels of E-Cad^+^ /N-Cad^+^ cells, and in many conditions, the expression levels of E-Cad and N-Cad are correlated rather than anticorrelated at the single cell level (Fig. S6). Even for established protocols that underwent cadherin switching, the degree of switching was highly variable (Fig. S1C). As these protocols were optimized to generate cell fates, but not to mimic in vivo levels of cadherin switching, and to our knowledge, quantitative measurements of cadherin expression in space and time have not been performed in vivo, it remains to be determined whether these differences are reflected in the embryo.

We modulated signaling and measured Snail1 and Cadherin expression to determine the relationship between them. On the first day, all pathways examined appeared to promote Snail and N-Cad expression, perhaps reflecting the activity of all these key pathways in initiating mesendodermal differentiation. The highest levels of Snail and N-Cad were found in MPS conditions which include activation of all four signals. In contrast, E-Cad retention was promoted by BMP, so that E-Cad expression was highest in PPS and lowest in APS. On the second day of differentiation, Activin and Wnt continued to promote Snail and N-Cad expression while the role of BMP changed so it repressed these factors. LM differentiation, which is performed in the presence of BMP, Wnt inhibition, and Nodal inhibition yielded the lowest levels of Snail and N-Cad.

Taken together our in vitro studies suggest two key points. First, that cells within the PS exhibit a wide window of potential which shifts gradually from anterior to poster. Second, that EMT and cadherin switching can be decoupled from cell fate acquisition and are regulated independently by signaling pathways. Future research will be needed to determine how these processes which can be regulated separately are tightly coupled during gastrulation in vivo.

## Methods

### Routine cell culture

All experiments were performed using the ESI017 hPSC line (obtained from ESI BIO, RRID: CVCL_B854, XX) or cell lines based on ESI017 (listed in Table S1). Cells were cultured in the chemically defined medium mTeSR1 or mTeSR Plus (Stemcell Technologies) in Matrigel-coated (Corning; 1:200 in PBS--, overnight at 4°C) 35 mm or 60mm culture dishes and kept at 37°C, 5% CO_2_. Cells were routinely passaged using dispase (Stemcell Technologies; 1:5 dilution in DMEM/F12 (VWR)) and checked for mycoplasma contamination with all negative results. In all experiments, cell passage number did not extend beyond 58. Single cell suspensions were prepared using accutase (Corning). ROCK-inhibitor Y-27672 (10uM; Stemcell Technologies) was used to maintain single cell status after seeding and before the start of treatment. 5ug/mL puromycin (Gibco) was used for selecting the CAAX:mCFP cell membrane marker.

### Differentiation

For differentiation, pluripotent hPSCs were prepared into a single cell suspension using accutase and seeded onto 96-well round µ-Plate or 18-well chambered µ-Slide (ibidi) with No. 1.5 polymer coverslip bottom, allowing high-resolution imaging. Before seeding, the plates or slides were coated with Matrigel (1:200 in PBS--, overnight at 4°C) or with Laminin-521 (Biolamina; 1:20 in PBS++, 2 hours at 37°C).

Definitive endoderm induction protocol was adapted from Loh *et al*. (2014). anterior and mid primitive streak, and paraxial and lateral mesoderm induction protocols was adapted from Loh *et al*. (2016). The adapted protocols are listed below:

First day treatment: After 12hrs or overnight, cells were differentiated into (1) definitive endoderm progenitor (or DE(a)) using 100 ng/mL Activin A + 2 uM CHIR99021 in Essential 6 medium (or E6; Gibco); (2) anterior primitive streak (APS) using 30 ng/mL Activin A + 20 ng/mL bFGF + 4 µM CHIR99021 in E6; (3) mid primitive streak (MPS) using 30 ng/mL Activin A + 20 ng/mL bFGF + 40 ng/mL Bmp4+ 6 µM CHIR99021 in E6; (4) posterior primitive streak (PPS) using 20 ng/mL bFGF + 40 ng/mL Bmp4+ 6 µM CHIR99021 in E6; (5) PS treatment without TGFβ reagents (noted as -TGFβ) using 20 ng/mL bFGF + 4 µM CHIR99021 in E6; or with reagents as described in the text. After 24-hour induction, cells were either further differentiated as described below, or fixed for immunostaining and imaging.

Second day treatment: After first day treatment, (1) DE(a) cells were further induced into definitive endoderm (or DE(b)) using 100 ng/mL Activin A +250 nM LDN-193189; (2) APS cells were further induced into paraxial mesoderm (PM) using 1 µM A-8301 + 20 ng/mL bFGF + 250 nM LDN-193189; (3) MPS cells were further induced into lateral mesoderm (LM) using 1 µM A-8301 + 30 ng/mL Bmp4 +4 µM IWP2. The first day and second day treatment were mixed to test the potency of each fate. After 48, cells were fixed for immunostaining and imaging

### Immunostaining

Cells were fixed with 4% PFA (Electron Microscopy Science) in PBS for 20 minutes at room temperature and washed twice with PBS. Fixed samples were then permeabilized and blocked with the block buffer (PBS with 0.1% Triton X100 (Sigma-Aldrich) + 3% donkey serum (Sigma-Aldrich)) for 1 hour at room temperature or overnight at 4 °C. Primary antibodies were diluted in the blocking buffer using ratios stated in Table S1, and applied to cells overnight at 4 °C. After three washes using PBST (PBS with 0.1% Tween 20 (Sigma Aldrich)), diluted secondary antibody (1:500 in the block buffer) with DAPI (1:5000) were applied to cells for 45-1hr at room temperature. After another three PBST washes, cells were maintained in PBS ready for imaging.

### Fixed cell imaging

Fixed cells were imaged on Olympus FV3000 laser scanning confocal microscope (LSM) with FV31S-SW version 2.4.1.198 software and a 20x, NA 0.75 objective or a 40x, NA 1.25 silicon oil objective. 5 to 8 positions were imaged for each condition.

### Live cell imaging

Doxycycline hyclate solution (Sigma-Aldrich) was added to medium 1 day before imaging to induce CAAX:mCerulean. Dual-reporter cells were seeded in µ-Plate or µ-Slide as described above and imaged on Olympus FV3000 LSM with environmental chamber (temperature at 37 °C, humidity at ∼50%, and CO_2_ at 5%), and a 40x, NA 1.25 silicon oil objective. 3 positions were imaged for each condition.

### Image analysis

All experiments were performed at least twice with consistent results. ‘n’ in the figure captions denotes the number of positions imaged per condition in the same experiment. In all cases, error bars represent standard error of the mean intensity.

For fixed cell imaging, maximum intensity projections were generated for each image using FIJI (Schindelin et al., 2012). Nuclear masks were created by pixel segmentation of the DAPI channel using ilastik (Berg et al., 2019), and subsequent analysis was performed using custom code written using MATLAB. Nuclear protein expression was measured as the mean nuclear intensity for each cell. Membrane protein expression was quantified using directional profiling, where fluorescence intensities were sampled along six equally spaced directions from the centroid of each nucleus. For each direction, the top 20% fluorescence intensity values within 25 pixels from the nucleus were averaged, and the final signal for each cell was calculated as the mean of all directional values.

For live cell imaging using ESI-017 E-Cad:mCitrine N-Cad:mCherry CAAX:mCerulane reporter cell line, pixel shift alignment was applied to align the three channels. A membrane mask and a non-membrane cell mask were generated based on the CAAX:mCerulane channel by pixel segmentation in ilastik. Background subtraction was then performed on the two membrane protein channels. Membrane protein expression was quantified as the ratio of mean membrane intensity to non-membrane cell intensity for each image.

MATLAB code is available in the following link: https://github.com/yz233333/CadherinSwitchingImageProcessing.

### Establishing ESI-017 E-Cad:mCitrine N-Cad:mCherry CAAX:mCerulane reporter cell line

The single reporter cell line ESI017 E-Cad:mCitrine was previously developed as stated in Liu *et al*. (2022). All plasmid constructions, DNA nucleofection, and antibiotic selection followed methods elaborated in Liu *et al*. (2022) but using different fluorescence proteins and targeting different genes.

### Plasmid constructions

1) N-Cad:mCherry: to insert mCherry into CDH2 (N-Cadherin) Exon16, (1) a Cas9 and sgRNA co-expression vector, px459 (Addgene) was modified to target CDH2 (N-cadherin) Exon16 (c), with sgRNA spacer sequence 5’-ACTGAACTTCAGGGTGAACT-3’; (2) A donor vector was created for homology directed DNA repair as follows: ∼400 bp of CDH2 pro-domain or mature domain sequence was inserted into the FloxP-neomycin resistant vector (AW-P216) as right/left homology arm, respectively, flanking the FloxP-neomycin expression cassette and mCherry. Cre Shine plasmid (Addgene #37404) was then used for excision of FloxP fragments after antibiotic-resistance selection.

2) CAAX:mCerulane: Restriction enzymes (BamHI and BsrGI) were used to digest the plasmids ePiggyBac based doxycycline inducible cell membrane marker mCherry-CAAX (Plasmid AW-P224 (Liu et al., 2022)) and pBSSK+-puroR-T2A-mCerulean (Plasmid AW-P130). The backbone from AW-P224 and the mCerulean insert from AW-P130 were gel extracted and ligated using T4 DNA ligase resulting in a replacement of the mCherry sequence in AW-P224 with mCerulean. The doxycycline inducible cell membrane marker mCerulean-CAAX plasmid (denoted AW-P235) was then transformed using NEB 10-beta competent E.coli C3019H.

### DNA nucleofection

All DNA nucleofections were performed using Amaxa P3 Primary Cell 4D-Nucleofector X Kit L (Lonza).

1)N-Cad:mCherry: insertion performed on ESI017 E-Cad:mCitrine cell using guide RNA, PX459-NCAD-sgRNA, and donors, mCherry loxp-N-loxp-G CN mat and NCad-mCherry-HDR. Cells were selected using G418 at 150ug/mL for one week. Then CRE Shine plasmid was nucleofected to remove antibiotics resistance sequence. The ESI-017 E-Cad:mCitrine-N-Cad:mCherry duo-reporter cell line were established

2)CAAX:mCerulane: insertion performed on ESI-017 E-Cad:mCitrine-N-Cad:mCherry and ESi-017 using: plasmid DNA of mCerulean-CAAX and ePB Helper (Gifted by ali Brivanlou (Rockefeller University)). Cells were selected using 5ug/mL puromycin. The ESI-017 E-Cad:mCitrine-N-Cad:mCherry-CAAX:mCerulane reporter cell line were established. Same insertion and selection were performed to build ESI-017 CAAX:mCerulane and ESI-017 E-Cad ^−/−^ CAAX:mCerulean cell lines.

### Establishing CDH1 (E-cadherin) knock out cell line

The E-Cad knock-out cell line with ESI-017 was established via CRISPR-Cas9 RNP (ribonucleoprotein) delivery (Park et al., 2022). Nucleofection was conducted using the Amaxa P3 Primary Cell 4D-Nucleofector X Kit S (Lonza), supplemented with SpCas9 Nuclease (IDT). sgRNA spacer (sequence 5’-G*U*G*AAUUUUGAAGAUUGCAC -3’) was designed using Sythego CRISPR Design Tool and produced by Sythego. The modified cells were sorted by single-cell flow cytometry using a SH800S Cell Sorter (Sony) with Cell Sorter software (Version 2.2.4.5150) into individual wells of 96-well plates. Sorted cells were maintained in mTeSR Plus medium supplemented with 1x CloneR (Stemcell Technologies). Single clones were established and then screened through Sanger sequencing (Fig. S10). Selected candidates were further verified through sequencing individual alleles with TOPO-cloning (Invitrogen).

### RNA extraction

hPSCs were differentiated for either one or two days as indicated. Cells were harvested using Accutase (Corning) and pelleted by centrifuging at 1000 rpm for 4 minutes. The supernatant was thoroughly removed by aspiration, and cell pellets were immediately disrupted in Lysis Solution. Then samples were snap-frozen in liquid nitrogen and stored at -80°C until RNA extraction. Total RNA was isolated using the RNAqueous™-Micro Total RNA Isolation Kit (Invitrogen) following the manufacturer’s protocol. Extracted RNA samples were transferred to fresh RNase-free tubes and stored at -20°C before sequencing. Bulk RNA sequencing was performed by Novogene Corporation Inc.

### RNA Sequencing Data Processing and Quantification

Raw RNA sequencing data were processed to obtain gene-level quantification of transcript abundance. Reads were aligned and quantified using Salmon (v1.10.2) (Patro et al., 2017) in quasi-mapping mode with sequence-specific and GC bias correction enabled to improve accuracy. A precomputed transcriptome index was used for quantification, based on the GCA_000001405.15 GRCh38_no_alt_analysis_set reference from NCBI (hg38 alias), generated using the selective alignment method (salmon_sa_index:default). Output files containing transcript-level abundance estimates were generated for each sample.

To integrate transcript annotations and facilitate downstream analysis, the quantification data were imported into R (v4.4.2), where transcript abundance estimates were summarized at the gene level. Metadata linking each sample to its experimental condition were incorporated to maintain experimental context. The resulting gene-level count matrix was exported as a CSV file for further analysis.

## Acknowledgements

We thank Sally Lowell, Guillaume Blin, Idse Heemskerk and members of the Warmflash lab for helpful discussions. We thank Elena Camacho Aguilar for adapting the image processing scripts for this work. We also thank Siqi Du for optimizing the CRISPR-Cas9 RNP delivery method for efficient knockout in hPSCs. This work was supported by grants to AW from NIH (R35GM149328 and R01HD112488), NSF (MCB-2135296), and Rice University (Edinburgh Rice award).

## References

Amack, J. D. (2021). Cellular dynamics of EMT: lessons from live in vivo imaging of embryonic development. Cell Communication and Signaling 19, 79.

Arnold, S. J., Hofmann, U. K., Bikoff, E. K. and Robertson, E. J. (2008). Pivotal roles for eomesodermin during axis formation,epithelium-to-mesenchyme transition and endoderm specification in the mouse. Development 135, 501–511.

Berg, S., Kutra, D., Kroeger, T., Straehle, C. N., Kausler, B. X., Haubold, C., Schiegg, M., Ales, J., Beier, T., Rudy, M., et al. (2019). ilastik: interactive machine learning for (bio)image analysis. Nat Methods 16, 1226–1232.

Bernardo, A. S., Faial, T., Gardner, L., Niakan, K. K., Ortmann, D., Senner, C. E., Callery, E. M., Trotter, M. W., Hemberger, M., Smith, J. C., et al. (2011). BRACHYURY and CDX2 Mediate BMP-Induced Differentiation of Human and Mouse Pluripotent Stem Cells into Embryonic and Extraembryonic Lineages. Cell Stem Cell 9, 144–155.

Burridge, P. W., Matsa, E., Shukla, P., Lin, Z. C., Churko, J. M., Ebert, A. D., Lan, F., Diecke, S., Huber, B., Mordwinkin, N. M., et al. (2014). Chemically defined generation of human cardiomyocytes. Nat Methods 11, 855–860.

Carver, E. A., Jiang, R., Lan, Y., Oram, K. F. and Gridley, T. (2001). The mouse snail gene encodes a key regulator of the epithelial-mesenchymal transition. Mol Cell Biol 21, 8184–8188.

Chen, B., Khan, H., Yu, Z., Yao, L., Freeburne, E., Jo, K., Johnson, C. and Heemskerk, I. (2024). Extended culture of 2D gastruloids to model human mesoderm development. bioRxiv 2024.03.21.585753.

Cheung, C., Bernardo, A. S., Trotter, M. W. B., Pedersen, R. A. and Sinha, S. (2012). Generation of human vascular smooth muscle subtypes provides insight into embryological origin-dependent disease susceptibility. Nat Biotechnol 30, 165–173.

Chhabra, S. and Warmflash, A. (2021). BMP-treated human embryonic stem cells transcriptionally resemble amnion cells in the monkey embryo. Biology Open 10, bio058617.

Chhabra, S., Liu, L., Goh, R., Kong, X. and Warmflash, A. (2019). Dissecting the dynamics of signaling events in the BMP, WNT, and NODAL cascade during self-organized fate patterning in human gastruloids. PLOS Biology 17, e3000498.

Ciruna, B. and Rossant, J. (2001). FGF Signaling Regulates Mesoderm Cell Fate Specification and Morphogenetic Movement at the Primitive Streak. Developmental Cell 1, 37–49.

Deng, C. X., Wynshaw-Boris, A., Shen, M. M., Daugherty, C., Ornitz, D. M. and Leder, P. (1994). Murine FGFR-1 is required for early postimplantation growth and axial organization. Genes Dev. 8, 3045–3057.

Gertow, K., Hirst, C. E., Yu, Q. C., Ng, E. S., Pereira, L. A., Davis, R. P., Stanley, E. G. and Elefanty, A. G. (2013). WNT3A Promotes Hematopoietic or Mesenchymal Differentiation from hESCs Depending on the Time of Exposure. Stem Cell Reports 1, 53–65.

Hsu, H.-T., Estarás, C., Huang, L. and Jones, K. A. (2018). Specifying the Anterior Primitive Streak by Modulating YAP1 Levels in Human Pluripotent Stem Cells. Stem Cell Reports 11, 1357–1364.

Huelsken, J., Vogel, R., Brinkmann, V., Erdmann, B., Birchmeier, C. and Birchmeier, W. (2000). Requirement for b-Catenin in Anterior-Posterior Axis Formation in Mice. The Journal of Cell Biology 148, 567–578.

Jo, K., Teague, S., Chen, B., Khan, H. A., Freeburne, E., Li, H., Li, B., Ran, R., Spence, J. R. and Heemskerk, I. (2022). Efficient differentiation of human primordial germ cells through geometric control reveals a key role for Nodal signaling. eLife 11, e72811.

Liu, P., Wakamiya, M., Shea, M. J., Albrecht, U., Behringer, R. R. and Bradley, A. (1999). Requirement for Wnt3 in vertebrate axis formation. Nat Genet 22, 361–365.

Liu, L., Nemashkalo, A., Rezende, L., Jung, J. Y., Chhabra, S., Guerra, M. C., Heemskerk, I. and Warmflash, A. (2022). Nodal is a short-range morphogen with activity that spreads through a relay mechanism in human gastruloids. Nat Commun 13, 497.

Loh, K. M., Ang, L. T., Zhang, J., Kumar, V., Ang, J., Auyeong, J. Q., Lee, K. L., Choo, S. H., Lim, C. Y. Y., Nichane, M., et al. (2014). Efficient Endoderm Induction from Human Pluripotent Stem Cells by Logically Directing Signals Controlling Lineage Bifurcations. Cell Stem Cell 14, 237–252.

Loh, K. M., Chen, A., Koh, P. W., Deng, T. Z., Sinha, R., Tsai, J. M., Barkal, A. A., Shen, K. Y., Jain, R., Morganti, R. M., et al. (2016). Mapping the Pairwise Choices Leading from Pluripotency to Human Bone, Heart, and Other Mesoderm Cell Types. Cell 166, 451–467.

Malaguti, M., Nistor, P. A., Blin, G., Pegg, A., Zhou, X. and Lowell, S. (2013). Bone morphogenic protein signalling suppresses differentiation of pluripotent cells by maintaining expression of E-Cadherin. eLife 2, e01197.

Martyn, I., Kanno, T. Y., Ruzo, A., Siggia, E. D. and Brivanlou, A. H. (2018). Self-organization of a human organizer by combined Wnt and Nodal signalling. Nature 558, 132–135.

Martyn, I., Brivanlou, A. H. and Siggia, E. D. (2019). A wave of WNT signaling balanced by secreted inhibitors controls primitive streak formation in micropattern colonies of human embryonic stem cells. Development 146, dev172791.

Mendjan, S., Mascetti, V. L., Ortmann, D., Ortiz, M., Karjosukarso, D. W., Ng, Y., Moreau, T. and Pedersen, R. A. (2014). NANOG and CDX2 Pattern Distinct Subtypes of Human Mesoderm during Exit from Pluripotency. Cell Stem Cell 15, 310–325.

Nakanishi, M., Kurisaki, A., Hayashi, Y., Warashina, M., Ishiura, S., Kusuda-Furue, M. and Asashima, M. (2009). Directed induction of anterior and posterior primitive streak by Wnt from embryonic stem cells cultured in a chemically defined serum-free medium. The FASEB Journal 23, 114–122.

Park, S. H., Cao, M., Pan, Y., Davis, T. H., Saxena, L., Deshmukh, H., Fu, Y., Treangen, T., Sheehan, V. A. and Bao, G. (2022). Comprehensive analysis and accurate quantification of unintended large gene modifications induced by CRISPR-Cas9 gene editing. Science Advances 8, eabo7676.

Patro, R., Duggal, G., Love, M. I., Irizarry, R. A. and Kingsford, C. (2017). Salmon provides fast and bias-aware quantification of transcript expression. Nat Methods 14, 417–419.

Patsch, C., Challet-Meylan, L., Thoma, E. C., Urich, E., Heckel, T., O’Sullivan, J. F., Grainger, S. J., Kapp, F. G., Sun, L., Christensen, K., et al. (2015). Generation of vascular endothelial and smooth muscle cells from human pluripotent stem cells. Nature cell biology 17, 994.

Punovuori, K., Migueles, R. P., Malaguti, M., Blin, G., Macleod, K. G., Carragher, N. O., Pieters, T., van Roy, F., Stemmler, M. P. and Lowell, S. (2019). N-cadherin stabilises neural identity by dampening anti-neural signals. Development 146, dev183269.

Schäfer, G., Narasimha, M., Vogelsang, E. and Leptin, M. (2014). Cadherin switching during the formation and differentiation of the Drosophila mesoderm – implications for epithelial-to-mesenchymal transitions. Development 141, e0907–e0907.

Scheibner, K., Schirge, S., Burtscher, I., Büttner, M., Sterr, M., Yang, D., Böttcher, A., Ansarullah Irmler, M., Beckers, J., et al. (2021). Epithelial cell plasticity drives endoderm formation during gastrulation. Nat Cell Biol 23, 692–703.

Schindelin, J., Arganda-Carreras, I., Frise, E., Kaynig, V., Longair, M., Pietzsch, T., Preibisch, S., Rueden, C., Saalfeld, S., Schmid, B., et al. (2012). Fiji: an open-source platform for biological-image analysis. Nat Methods 9, 676–682.

Smith, D. E., Amo, F. F. D. and Gridley, T. (1992). Isolation of Sna, a mouse gene homologous to the Drosophila genes snail and escargot: its expression pattern suggests multiple roles during postimplantation development. Development 116, 1033–1039.

Umeda, K., Zhao, J., Simmons, P., Stanley, E., Elefanty, A. and Nakayama, N. (2012). Human chondrogenic paraxial mesoderm, directed specification and prospective isolation from pluripotent stem cells. Sci Rep 2, 455.

Yamaguchi, T. P., Harpal, K., Henkemeyer, M. and Rossant, J. (1994). fgfr-1 is required for embryonic growth and mesodermal patterning during mouse gastrulation. Genes Dev. 8, 3032–3044.

Yang, J., Antin, P., Berx, G., Blanpain, C., Brabletz, T., Bronner, M., Campbell, K., Cano, A., Casanova, J., Christofori, G., et al. (2020). Guidelines and definitions for research on epithelial– mesenchymal transition. Nat Rev Mol Cell Biol 21, 341–352.

