## Supplementary figures for "Dependence of cell fate potential and cadherin switching on primitive streak coordinate during differentiation of human pluripotent stem cells"

### Supporting information

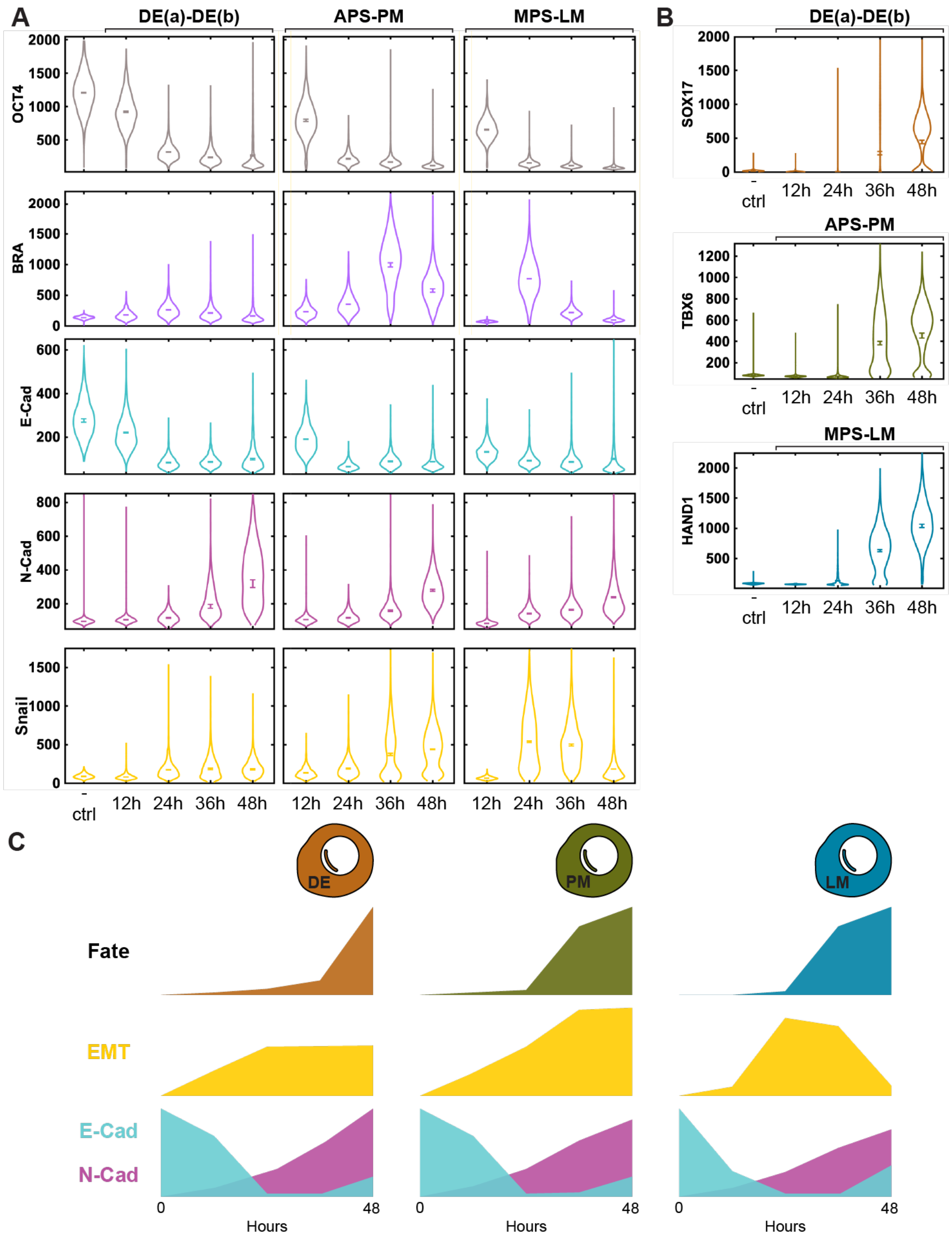


**Supplementary Figure 1. Dynamics of fate commitment in LM, PM, and DE differentiation.** Quantification of nuclear and membrane marker expression of DE(a)-DE(b), APS-PM, and MPS-LM treated cells at 12hr, 24hr, 36hr, and 48hr after induction start, using confocal microscopy. - ctrl: negative control maintained in mTeSR Plus. **A.** OCT4: pluripotency marker. BRA: PS transient marker; E-Cad: E-Cadherin. N-Cad: N-Cadherin. Snail: EMT marker. **B.** SOX17: DE fate marker. TBX6: PM fate marker. HAND1: LM fate marker. (N = 7 images per condition.). **C**. Schematic comparing timing of fate marker expression, cadherin switching, and EMT during differentiation protocols.

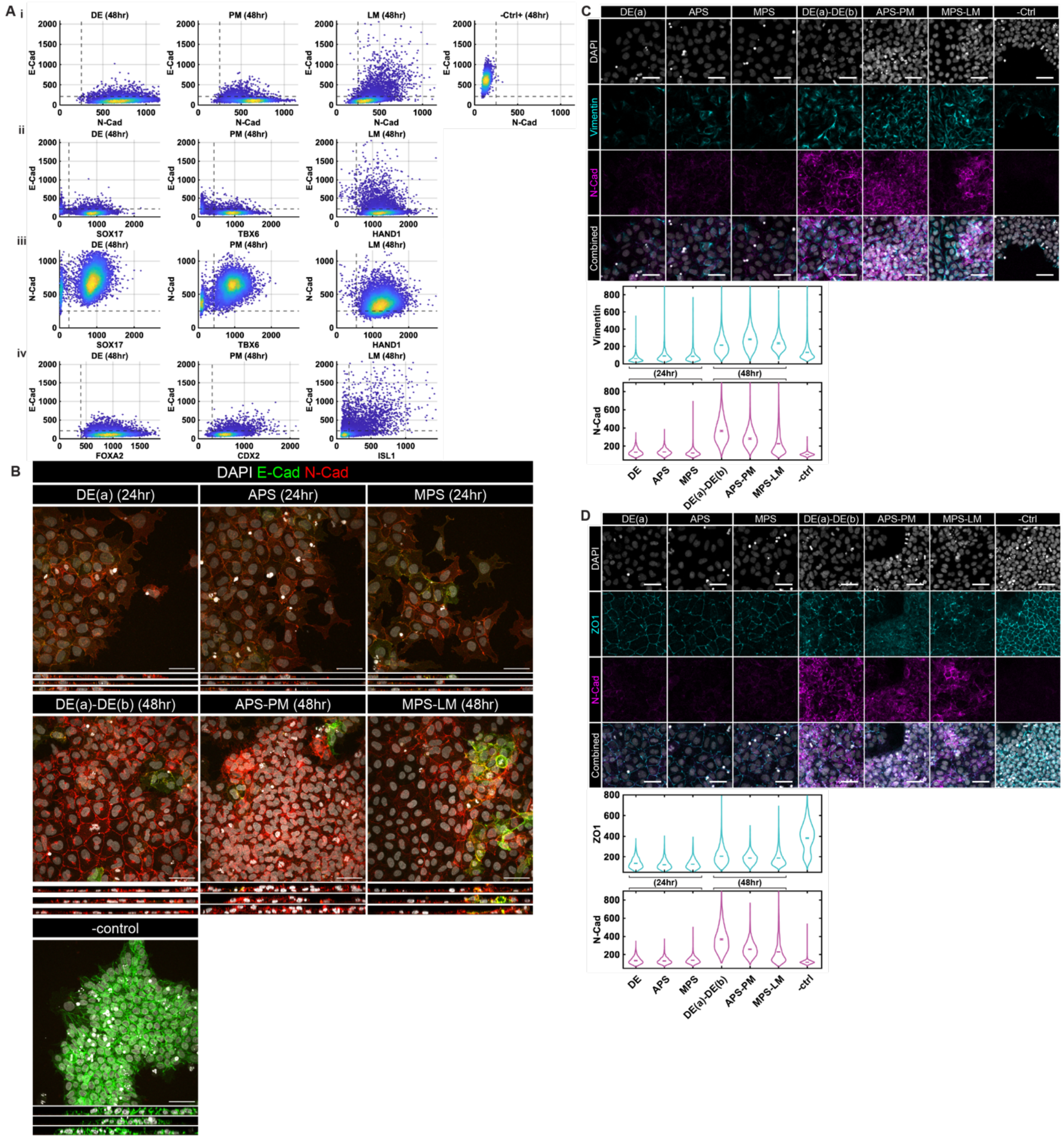


**Supplementary Figure 2. Correlation between cadherins and fate commitment in mesendodermal differentiation. A.** Co-expression scatter plots of E-Cad and N-Cad (**i**), E-Cad and fate markers (**ii** and **iv**), and N-Cad and fate markers (**iii**) for DE, PM and LM cells after 48hr treatment. – Ctrl: negative control that is maintained in mTeSR Plus. SOX17 and FOXA2: DE fate marker. TBX6 and CDX2: PM fate marker. HAND1 and ISL1: LM fate marker. (N = 6 images per condition.) **B**. Example confocal immunofluorescent images of high magnification (40x) and orthogonal view of DE(a), APS, MPS, DE(a)-DE(b), APS-PM, MPS-LM, and epithelial control cells (maintained in mTeSR Plus) immunostained with E-Cad, N-Cad, and DAPI. Scale Bars: 50 μm. **C and D**. Example confocal immunofluorescent images and

and quantification based on fluorescent intensity of Vimentin (**C**) and ZO-1 (**D**) for DE(a), APS, MPS, DE(a)-DE(b), APS-PM, MPS-LM, and epithelial control cells (maintained in mTeSR Plus). Scale Bars: 50 μm. (N = 7 images per condition.)

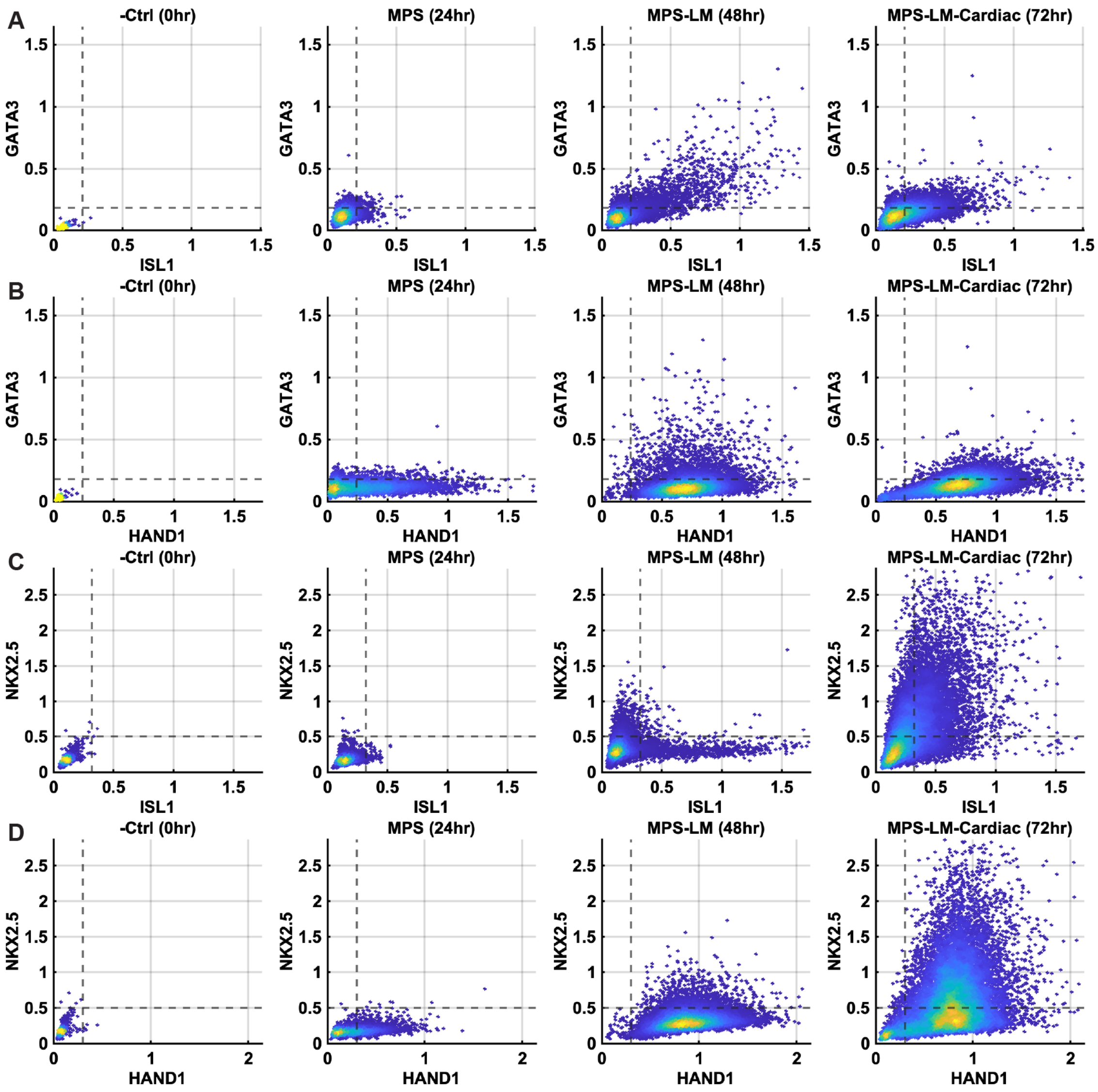


**Supplementary Figure 3. Cells in LM conditions contain a subset of amnion cells and are competent to generate cardiac progenitors.** Co-expression scatter plots of GATA3 and ISL1 (**A**), GATA3 and HAND1 (**B**), NKX2.5 and ISL1 (**C**), and NKX2.5 and HAND1 (**D**) for cells after 0hr, 24hr, 48hr, and 72hr MPS-LM-Cardiac treatment (0-24hr in MPS treatment; 24-48hr in LM treatment; 48-72hr in cardiac treatment). – Ctrl: negative control that is maintained in mTeSR Plus, fixed at 0hr. GATA3: amnion marker. NKX2.5: cardiac marker. HAND1 and ISL1: LM fate marker. (N=6 images per condition.)

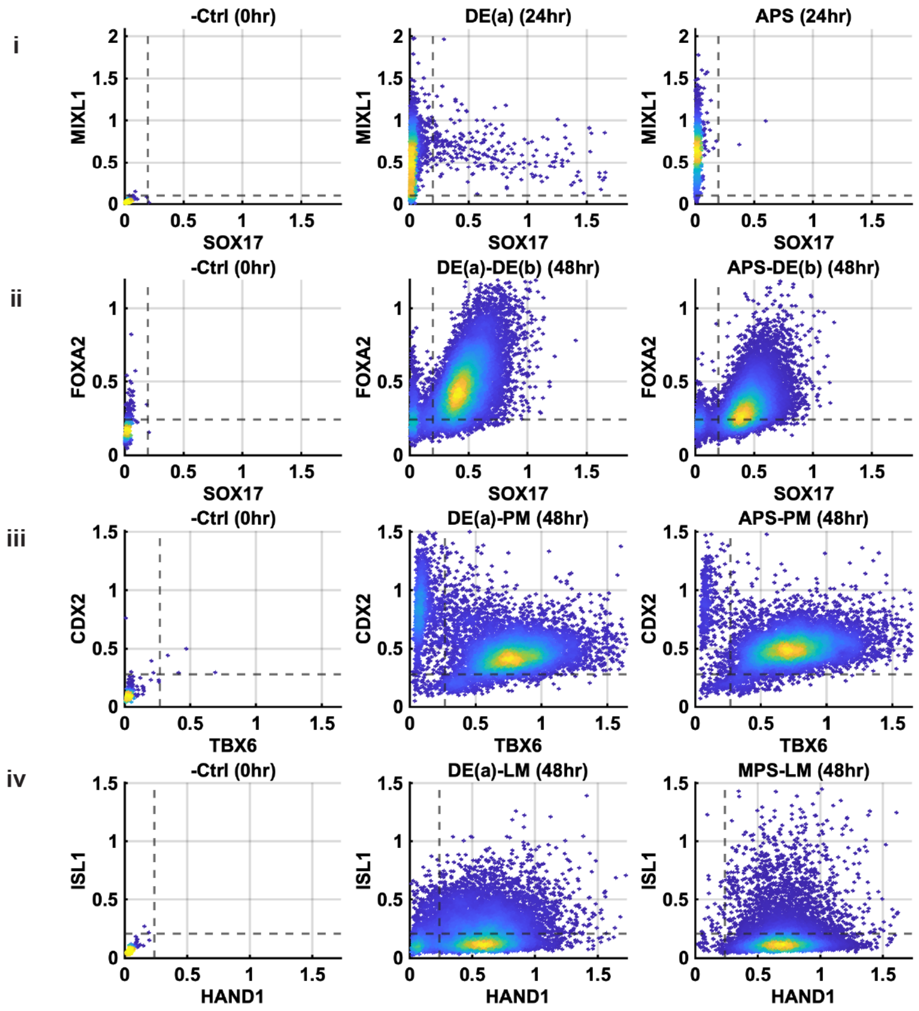


**Supplementary Figure 4. Cells treated with DE(a) are competent to generate DE, PM, and LM.** Co-expression scatter plots of (i) MIXL1 and SOX17 for DE(a), and APS cells after 24hr induction; (ii) FOXA2 and SOX17 for DE(a)-DE(b), and APS-DE(b) cells after 48hr induction; (iii) CDX2 and TBX6 for DE(a)-PM, and APS-PM cells after 48hr induction; (iv) ISL1 and HAND1 for DE(a)-LM, and MPS-LM cells after 48hr induction. (N = 6 images per condition.)

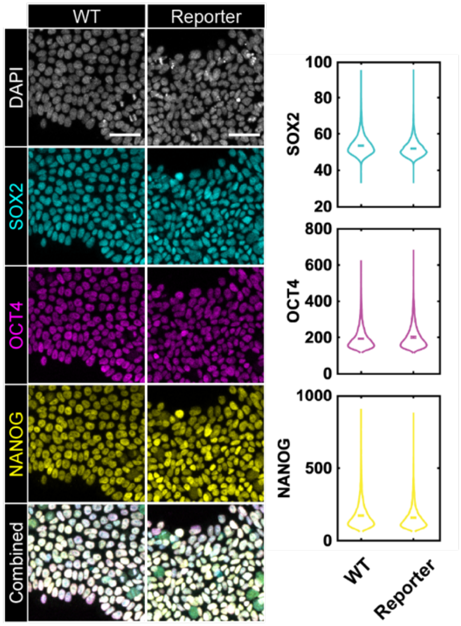


**Supplementary Figure 5. Validation of ESI017 CAAX:mCerulean cell line.**

Example confocal immunofluorescent images and quantification based on fluorescent intensity of pluripotency markers, SOX2, OCT4, and NANOG for ESI017 wildtype (WT), and CAAX:mCerulean (maintained in mTeSR Plus). DAPI: nuclear marker. Scale Bars: 50 µm. (N = 7 images per condition.)

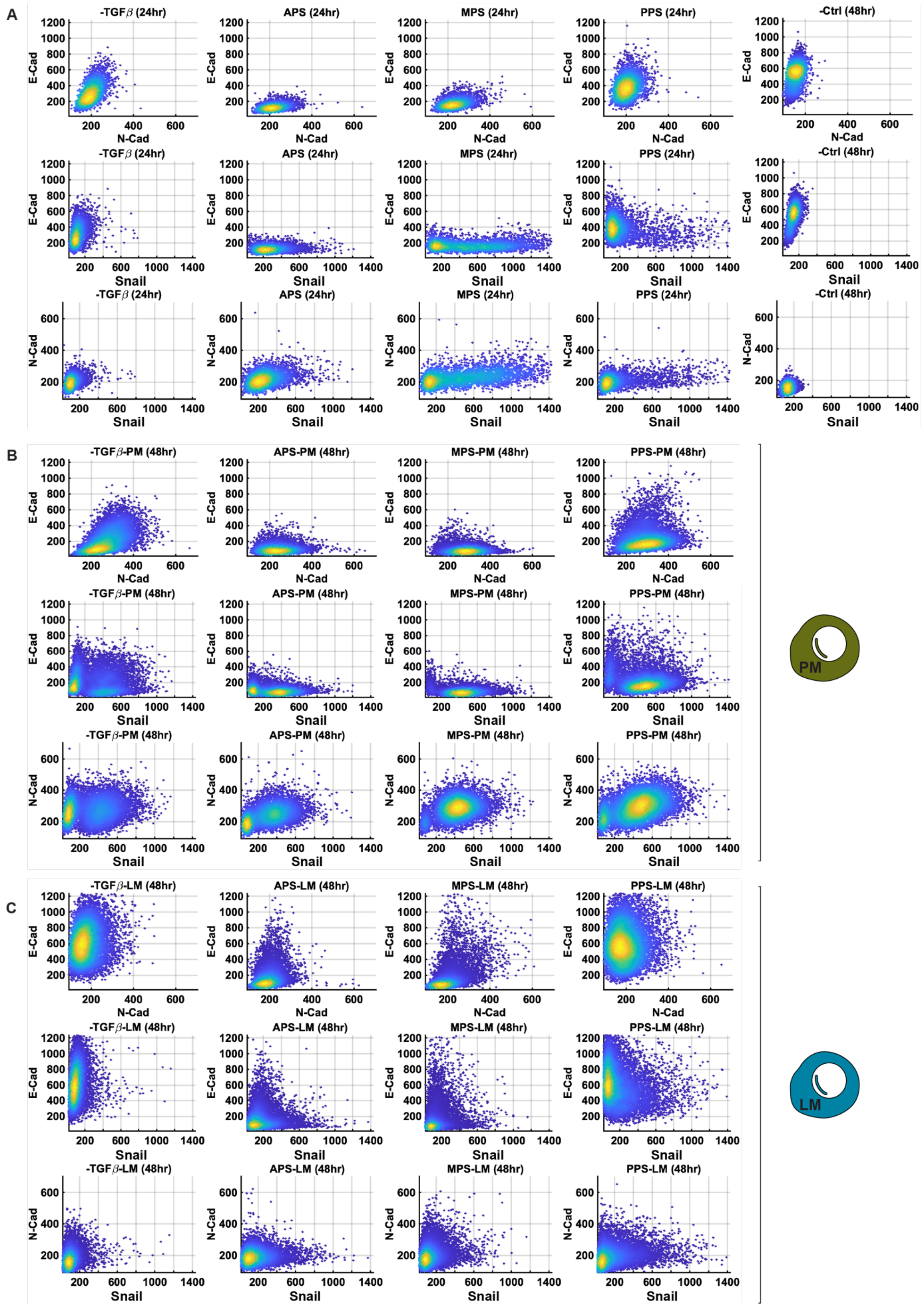
A
-TGFβ (24hr)
APS (24hr)
MPS (24hr)
PPS (24hr)
-Ctrl (48hr)
E-Cad
N-Cad
Snail
B
-TGFβ-PM (48hr)
APS-PM (48hr)
MPS-PM (48hr)
PPS-PM (48hr)
PM
C
-TGFβ-LM (48hr)
APS-LM (48hr)
MPS-LM (48hr)
PPS-LM (48hr)
LM

**Supplementary Figure 6. Snail1, E-Cad, and N-cad expression are not correlated.** Co-expression scatter plots of E-Cad and N-Cad, E-Cad and Snail, and N-Cad and Snail for cells treated by -TGF$\beta$, APS, MPS or PPS and fixed by 24hr, along with negative control group (-Ctrl, maintained in mTeSR Plus and fixed at 0hr) (**A**); and also, for the four PS treated groups further induced towards PM (**B**) or LM (**C**). (N = 8 images per condition.)

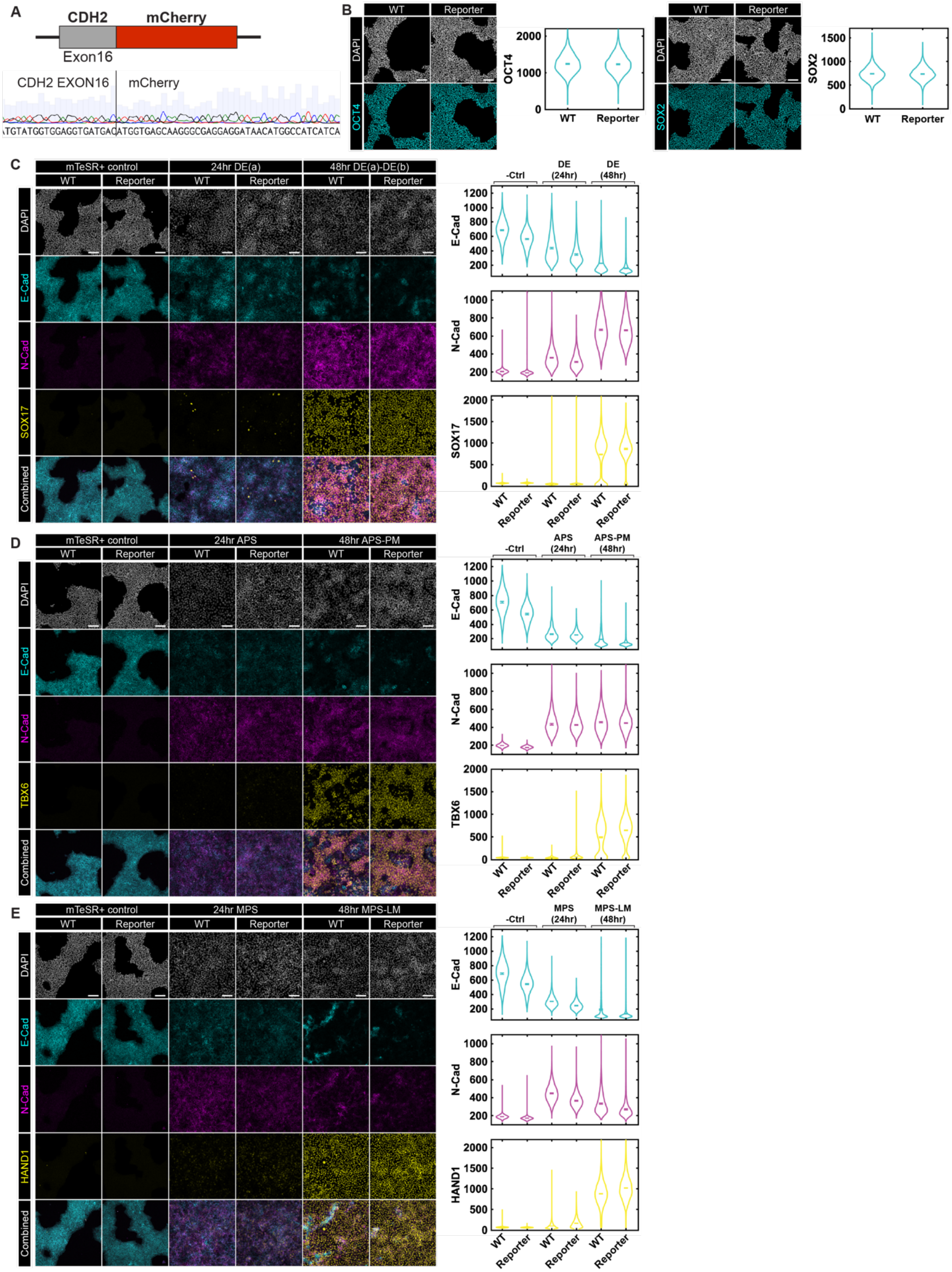


**Supplementary Figure 7. Characterization of the E-Cad/N-Cad dual reporter line. A**. Sequencing result of modified CDH2 locus showing connection of CDH2 Exon 16 and mCherry coding sequence. The base cell line is ESI-017 CDH1:mCitrine. CAAX-mCerulean was also inserted to make ESI017 CDH1:mCitrine-CDH2:mCherry-CAAX:mCeruleanReporter cell line. **B**. Pluripotency validation of ESI017-CDH1:mCitrine-CDH2:mCherry-CAAX:mCeruleancell line. Example images and quantification based on fluorescent intensity of pluripotency markers, OCT4, and SOX2, for ESI017 wildtype (WT), and ESI017-CDH1:mCitrine-CDH2:mCherry-CAAX:mCerulean(maintained in mTeSR Plus). Scale Bars: 50 μm. **C**, **D** and **E**. Example confocal immunofluorescent images and quantification based on fluorescent intensity of E-Cad, N-Cad and fate markersfor ESI017-CDH1:mCitrine-CDH2:mCherry-CAAX:mCeruleanReporter cell line and wild type (WT) ESI0-17 underwent DE(a)-DE(b) (SOX17) (**C**), APS-PM (TBX6) (**D**), and MPS-LM (HAND1) (**E**) treatment, respectively. DAPI: nuclear marker. Scale Bars: 50 μm. (N = 7 images per condition.)

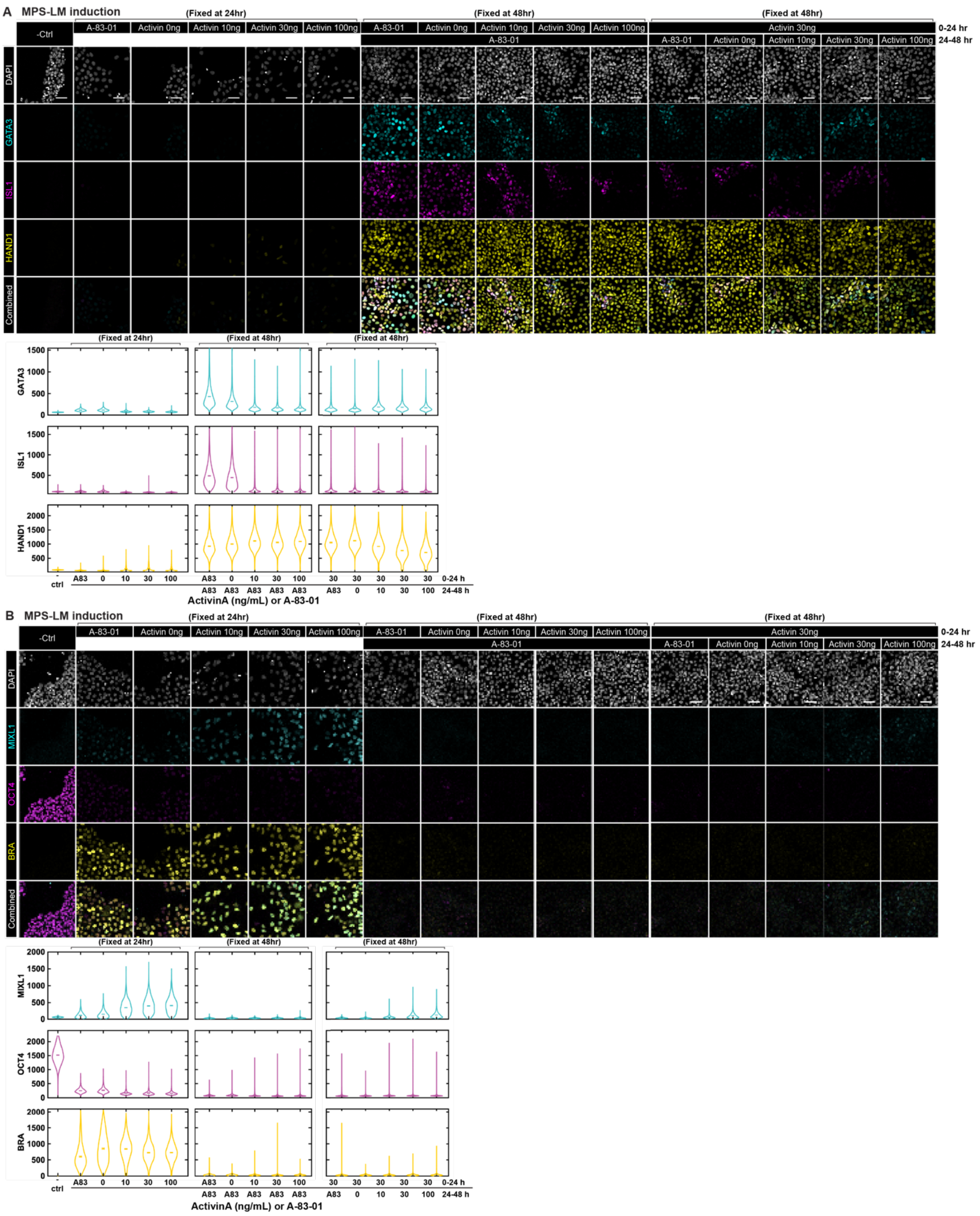


**Supplementary Figure 8. Effects of varying Activin signaling levels on LM fate commitment.**
Example confocal immunofluorescent images and quantification based on fluorescent intensity of GATA3, ISL1, and HAND1 (**A**), and MIXL1, OCT4 and BRA (**B**) for cells underwent MPS induction with A-83-01 (1 μM) or various Activin A doses (0, 10, 30, and 100 ng/mL) and fixed by 24hr; for all the MPS induced cells

continued with LM treatment (A-83-01 1 μM) and fixed by 48hr; and for cells underwent original MPS treatment (Activin A at 30 ng/mL), and switched to LM treatment with A-83-01 (1 μM) or various Activin A doses (0, 10, 30, and 100 ng/mL) and fixed by 48hr. (N = 7 images per condition.)

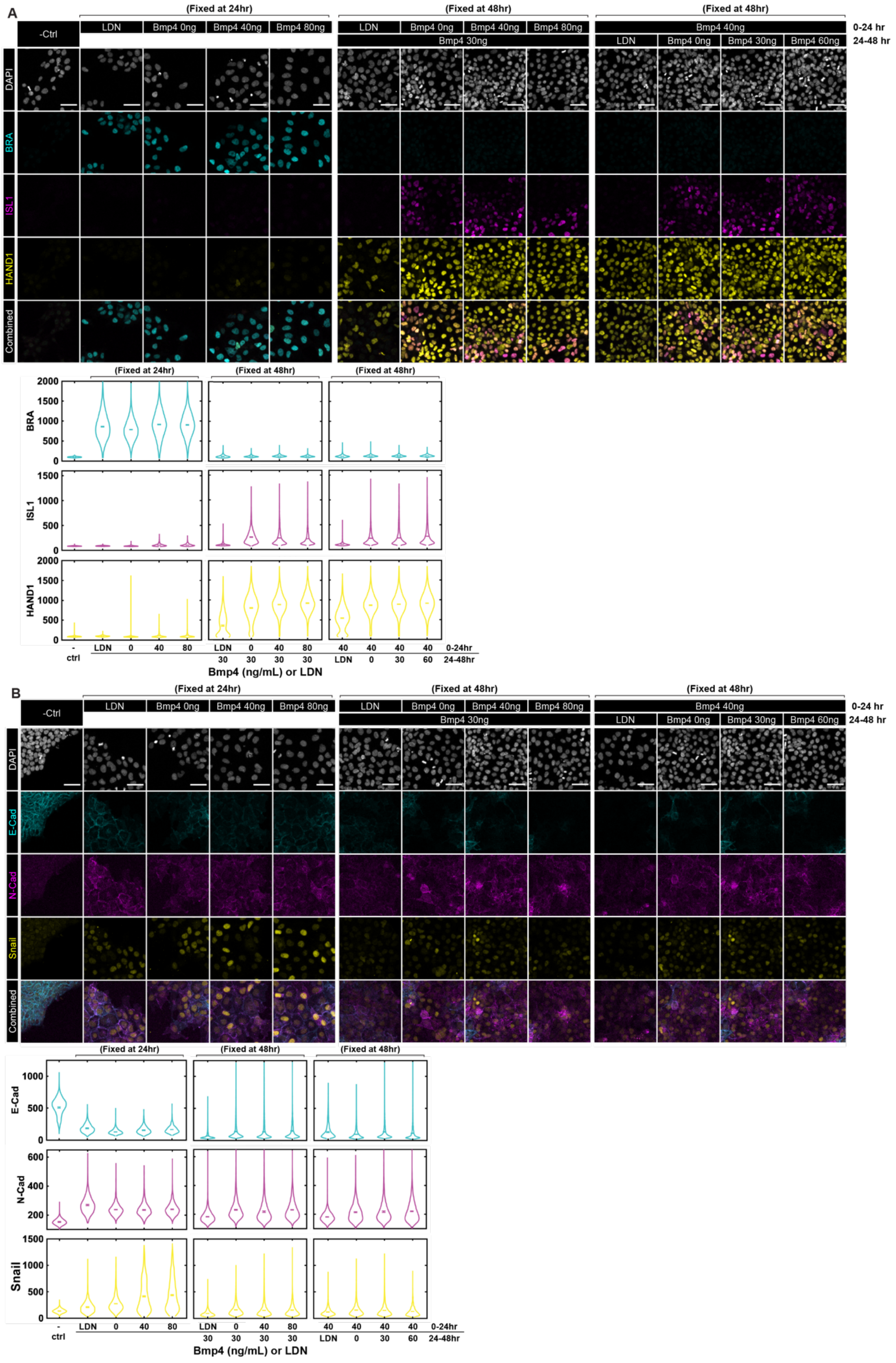
A
(Fixed at 24hr)
(Fixed at 48hr)
(Fixed at 48hr)
-Ctrl
LDN
Bmp4 0ng
Bmp4 40ng
Bmp4 80ng
Bmp4 30ng
Bmp4 40ng
Bmp4 30ng
Bmp4 60ng
0-24 hr
24-48 hr
DAPI
BRA
ISL1
HAND1
Combined
Bmp4 (ng/mL) or LDN
0-24hr
24-48hr
ctrl
B
E-Cad
N-Cad
Snail

**Supplementary Figure 9. Effects of varying BMP signaling levels on EMT and cadherin switching.** Example confocal immunofluorescent images and quantification based on fluorescent intensity of BRA, ISL1, and HAND1 (**A**), and E-CAD, N-Cad and Snail (**B**) for cells underwent MPS induction with LDN or various Bmp4 doses (0, 40, and 80 ng/mL) and fixed by 24hr; for all the MPS induced cells continued with LM treatment (Bmp4 at 30 ng/mL) and fixed by 48hr; and for cells underwent original MPS treatment (Bmp4 at 40 ng/mL), and switched to LM treatment with LDN or various Bmp4 doses (0, 30, and 60 ng/mL) and fixed by 48hr. (N = 8 images per condition.)

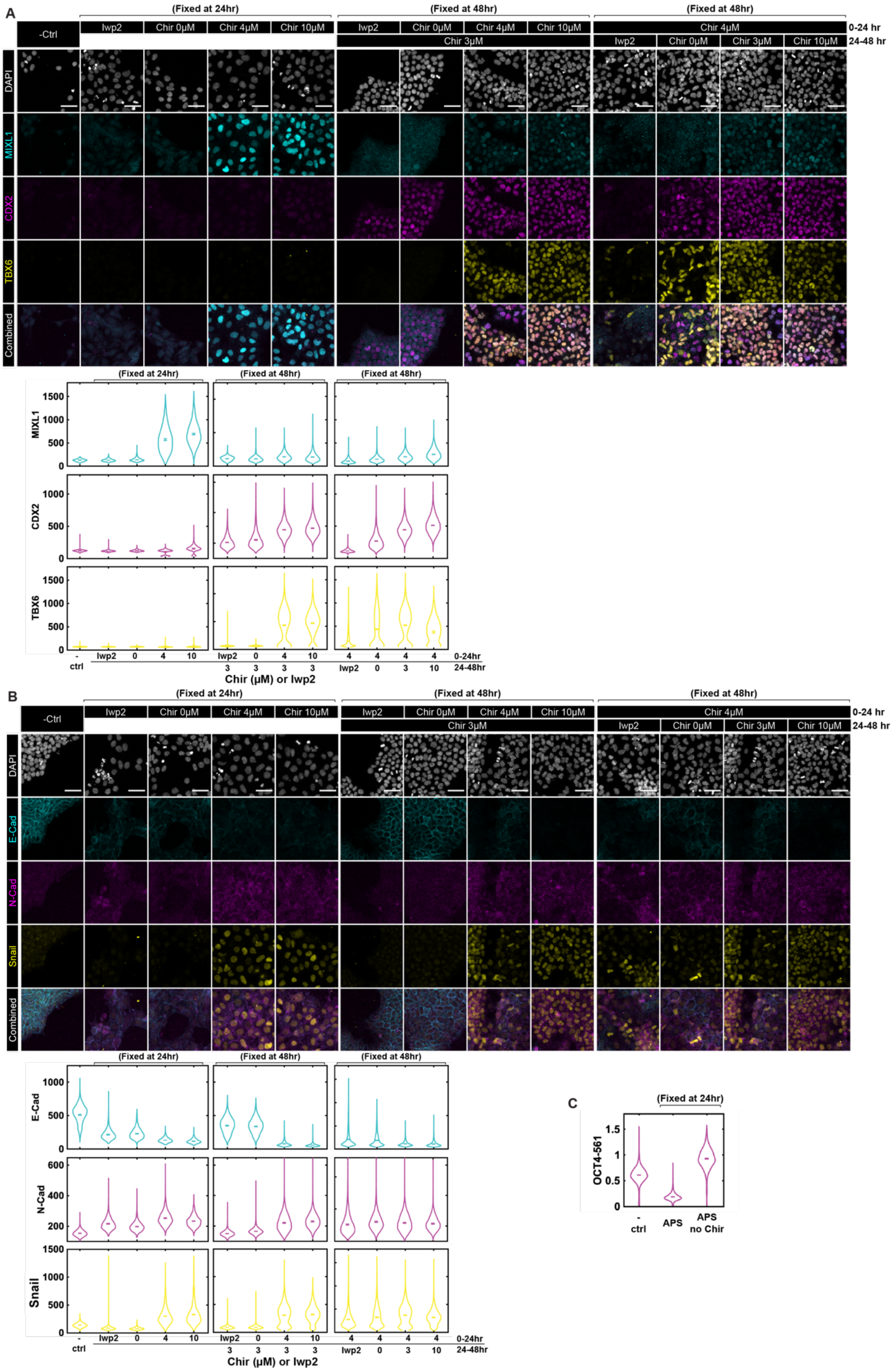
A
(Fixed at 24hr)
(Fixed at 48hr)
(Fixed at 48hr)
-Ctrl
Iwp2
Chir 0μM
Chir 4μM
Chir 10μM
Chir 3μM
0-24 hr
24-48 hr
DAPI
MIXL1
CDX2
TBX6
Combined
Chir (μM) or Iwp2
ctrl
0-24hr
24-48hr
B
E-Cad
N-Cad
Snail
C
OCT4-561
APS
APS no Chir

75

**Supplementary Figure 10. Effects of varying WNT signaling levels on EMT and cadherin switching.** Example confocal immunofluorescent images and quantification based on fluorescent intensity of MIXL1, CDX2, and TBX6 (**A**), and E-CAD, N-Cad and Snail (**B**) for cells underwent APS induction with IWP2 or various CHIR doses (0, 4, and 10 μM) and fixed by 24hr; for all the APS induced cells continued with PM treatment (CHIR at 3 μM) and fixed by 48hr; and for cells underwent original APS treatment (CHIR at 4 μM), and switched to PM treatment with IWP2 or various CHIR doses (0, 3, and 10 μM) and fixed by 48hr. **C**. Quantification based on inflorescent intensity of pluripotency marker, OCT4, for cells treated using mTeSR Plus (- Ctrl), APS, or APS with no CHIR. (N = 8 images per condition.)

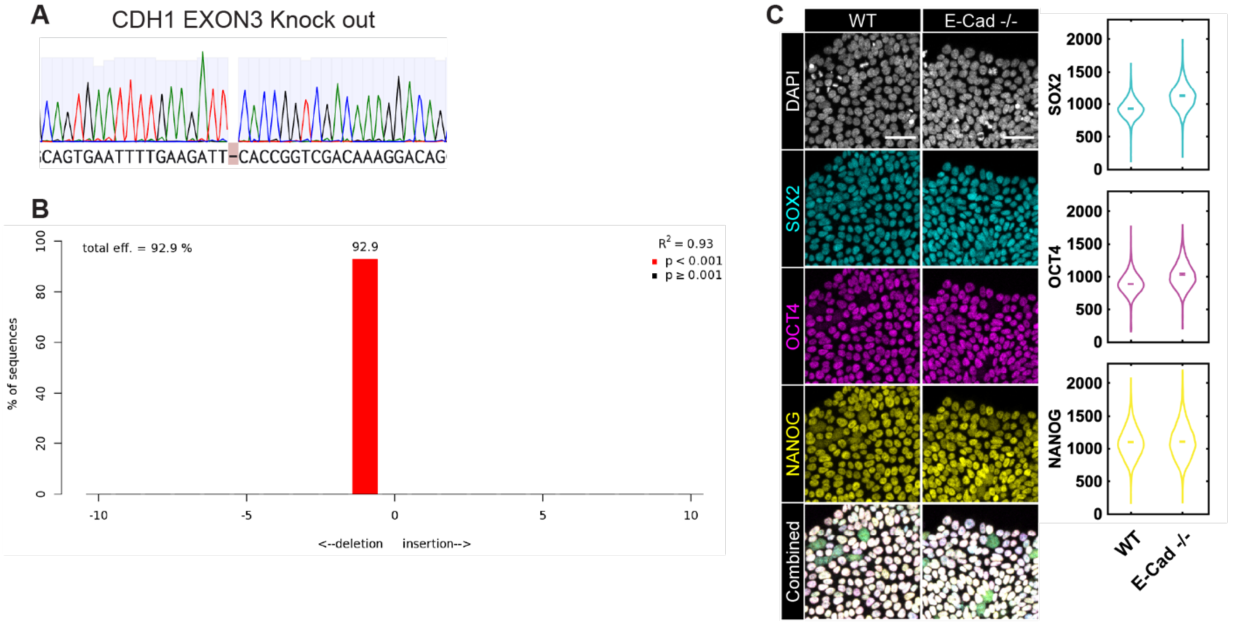


**Supplementary Figure 11. Validation of CDH1 (E-Cadherin) knock-out. A**. ESI017-CDH1 $^{-/-}$ was verified by sequencing, where the same deletion of 1 bp causing open reading frame (ORF) shifts for both alleles. **B**. A comprehensive profile of all insertions and deletions in the ESI017-CDH1 $^{-/-}$ sample by TIDE analysis (Brinkman et al., 2014). **C**. Pluripotency validation of ESI017- CDH1 $^{-/-}$ cell line. Example confocal immunofluorescent images and quantification based on fluorescent intensity of pluripotency markers, SOX2, OCT4, and NANOG for ESI017 wildtype (WT), and CDH1 $^{-/-}$ (maintained in mTeSR Plus). DAPI: nuclear marker. Scale Bars: 50 μm. (N = 7 images per condition.)

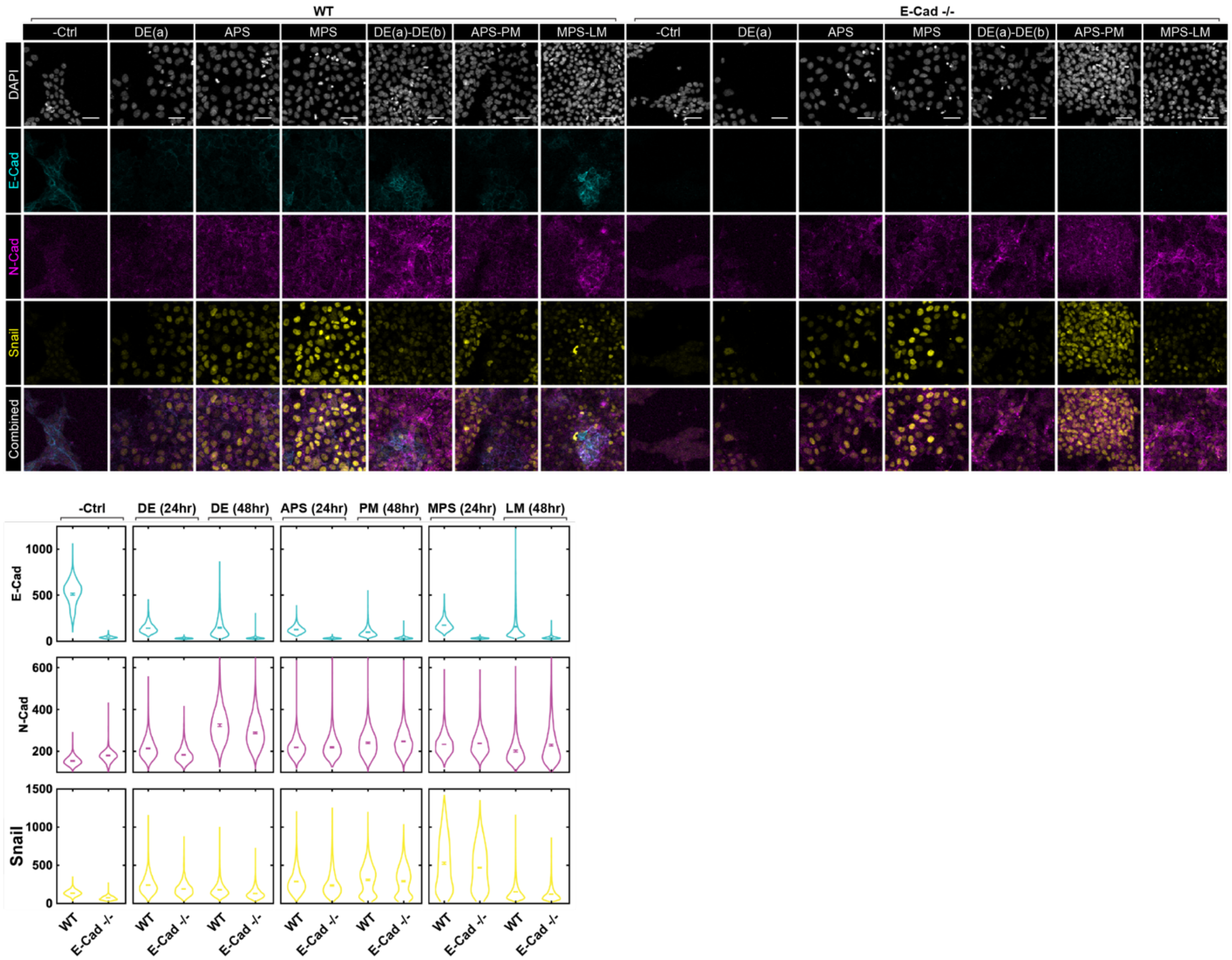


**Supplementary Figure 12. Effects of loss of E-CAD on N-CAD and Snail expression during differentiation.** Example confocal immunofluorescent images and quantification based on fluorescent intensity of E-Cad, N-Cad, and Snail for ESI017 wildtype (WT), and CDH1 $^{-/-}$ induced towards DE(a), APS, and MPS (fixed by 24hr), and further induced towards DE(b), PM and LM. DAPI: nuclear marker. Scale Bars: 50 µm. (N = 8 images per condition.)

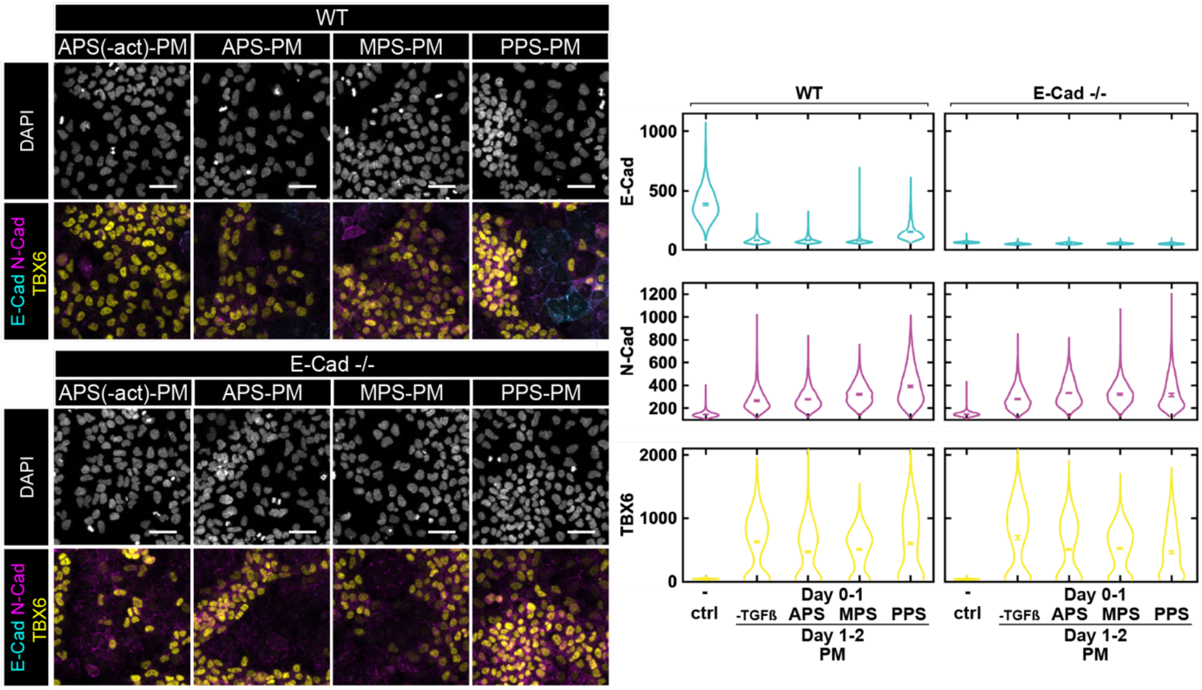


**Supplementary Figure 13. Loss of E-CAD did not affect fate outcomes during PM differentiation**

Example confocal microscopic images and quantification of E-Cad, N-Cad, and fate markers for WT and E-Cad $^{-/-}$ cells treated by - TGFβ, APS, MPS or PPS (0 – 24 hr), further induced to PM (24 – 48 hr; fate marker – TBX6), and fixed at 48 hr. Scale Bars: 50 µm. – Ctrl: negative control that is maintained in mTeSR Plus. (N=5 images per condition.)

**Supplementary Table 1. Key Resource Table.**

| REAGENT or RESOURCE | SOURCE | IDENTIFIER |
|---|---|---|
| ***Multi-well plate for imaging*** | | |
| μ-Plate 96 Well Round Black (with No. 1.5 polymer coverslip bottom) | Ibidi | Cat# 89606 |
| μ-Slide 18 Well (with No. 1.5 polymer coverslip bottom) | Ibidi | Cat# 81816 |
| ***Chemicals, Peptide, and Recombinant Proteins*** | | |
| mTeSR1 | Stemcell Technologies | Cat# 85875 |
| mTeSR Plus | Stemcell Technologies | Cat# 100-0276 |
| Matrigel | Corning | Cat# 354277 |
| Dulbecco's PBS without calcium and magnesium | Caisson Labs | Cat# PBL01-6X500ML |
| Dispase | Stemcell Technologies | Cat# NC9886504 |
| DMEM/F12 | VWR | Cat# 45000-344 |
| Accutase | Innovative Cell Technologies | Cat# NC9839010 |
| Y-27632 dihydrochloride (ROCK inhibitor) | Tocris | Cat# 1254/10 |
| Human recombinant laminin-521 protein | Biolamina | Cat# R021599/X0086842 |
| Dulbecco's PBS with calcium and magnesium | Caisson Labs | Cat# PBL02-6X500ML |
| Essential 6 medium | Gibco | Cat# A15165-01 |
| Recombinant Human/Mouse/Rat Activin A Protein | R&D Systems | Cat# 338-AC |
| CHIR99021 | Medchemexpress LLC | Cat# 50-187-1892 |
| Human FGF-basic (FGF-2/bFGF) (154 aa), Animal-Free Recombinant Protein | Gibco | Cat# AF-100-18B-100UG |
| Recombinant Human BMP-4 Protein | R&D Systems | Cat# 314-BP-050 |
| LDN-193189 | Stemolecule | Cat# 04-0074-02 |
| A 83-01 | Stemcell Technologies | Cat# 72022 |
| IWP-2 | Stemgent | Cat# 04-0034 |

| | | |
|---|---|---|
| Paraformaldehyde Aqueous Solution (PFA) | Electron Microscopy Science | Cat# 15710 |
| Triton X-100 | Sigma-Aldrich | Cat# 1001843780 |
| Donkey Serum | Sigma-Aldrich | Cat# S30-100ML |
| Tween 20 | Sigma-Aldrich | Cat# P1379 |
| DAPI (4,6-diamidino-2-phenylindole, dihydrochloride) | Invitrogen | Cat# D1306 |
| Clone R | Stemcell Technologies | Cat# 05889 |
| Amaxa P3 Primary Cell 4D-Nucleofector X kit L | Lonza | Cat# V4XP-3012 |
| Amaxa P3 Primary Cell 4D-Nucleofector X kit S | Lonza | Cat# V4XP-3032 |
| TOPO™ TA Cloning™ Kit for Sequencing, without competent cells | Invitrogen | Cat# 450030 |
| OneTaq Hot Start | NEB | Cat# M0489S |
| SpCas9 Nuclease | IDT | Cat# 1081061 |
| Puromycin Dihydrochloride | Gibco | Cat# A1113803 |
| Doxycycline hyclate | Sigma-Aldrich | Cat# D9891-1G |
| ***Antibodies*** | | |
| Rabbit Anti-E-Cad (1:300) | Cell Signaling Technology | Cat# 3195S |
| Mouse Anti-N-Cad (1:300) | Cell Signaling Technology | Cat# 14215S |
| Goat Anti-Snail (1:200) | R&D Systems | Cat# AF3639 |
| Rabbit Anti-BRACHYURY (1:400) | R&D Systems | Cat# MAB20851 |
| Mouse Anti-OCT3/4 (1:200) | BD Biosciences | Cat# 611203 |
| Mouse Anti-FOXA2 (1:200) | BD Biosciences | Cat# 561580 |
| Goat Anti-SOX17 (1:200) | R&D Systems | Cat# AF1924 |
| Goat Anti-BRACHYURY (1:300) | R&D Systems | Cat# AF2085 |
| Rabbit Anti-GATA3 (1:300) | Invitrogen | Cat# PA1-101 |
| Mouse Anti-ISL1 (1:75) | Developmental Studies Hybridoma Bank | Antibody Registry ID# AB_2314683 |
| Goat Anti-HAND1 (1:200) | R&D Systems | Cat# AF3168 |
| Rabbit Anti-MIXL1 (1:300) | ABclonal | Cat# A17223 |
| Mouse Anti-CDX2 (1:100) | Developmental Studies Hybridoma Bank | Antibody Registry ID# AB_2618482 |
| Goat Anti-TBX6 (1:200) | R&D Systems | Cat# AF4744-SP |

| | | |
|---|---|---|
| Rabbit Anti-SOX2 (1:200) | Cell Signaling Technologies | Cat# 5024S |
| Goat Anti-Nanog (1:400) | R&D Systems | Cat# AF1997 |
| Goat Anti-GFP (1:400) | Rockland | Cat# 600-101-215s |
| Goat Anti-FOXF1 (1:200) | R&D Systems | Cat# AF4798 |
| Rabbit Anti-Vimentin (1:300) | Cell Signaling Technology | Cat# 5741 |
| Rabbit Anti-ZO-1 (1:300) | Cell Signaling Technology | Cat# 5406S |
| Mouse Anti-EpCam (1:300) | Cell Signaling Technology | Cat# 2929T |
| Donkey anti-Rabbit IgG (H+L) Highly Cross-Adsorbed Secondary Antibody, Alexa Fluor™ 488 | Invitrogen | Cat# A-21206 |
| Donkey anti-Mouse IgG (H+L) Highly Cross-Adsorbed Secondary Antibody, Alexa Fluor™ 555 | Invitrogen | Cat# A-31570 |
| Donkey anti-Goat IgG (H+L) Cross-Adsorbed Secondary Antibody, Alexa Fluor™ 647 | Invitrogen | Cat# A-21447 |
| RNAqueous™-Micro Total RNA Isolation Kit | Invitrogen | Cat# AM1931 |
| ***Experimental Models: Cell Lines*** | | |
| ESI-017 | ESI BIO | RRID:CVCL_B854 |
| ESI-017 CAAX-mCFP (dox inducible) | | |
| ESI-017 E-Cad:mCitrine - N-Cad:mCherry-CAAX:mCerulean(dox inducible) | | |
| ESI-017 E-Cad $^{-/-}$ | | |
| ***Software and Algorithms*** | | |
| MATLAB | | https://www.mathworks.com/products/matlab.html |
| ilastik | (Berg et al., 2019) | http://ilastik.org/ |
| FIJI | (Schindelin et al., 2012) | https://fiji.sc/ |
| TIDE | (Brinkman et al., 2014) | http://shinyapps.datacurators.nl/tide/ |
| Salmon | (Patro et al., 2017) | |

**Supplementary Table 2. Primers for genomic DNA PCR**

| Target Gene | Forward | Reverse |
|---|---|---|
| CDH2 | 5' AAGGCAGTGGCTCCACTGC 3' | 5' ACTGATATTCCCTCTGAGCCC 3' |
| CDH2-mCherry-HDR | 5' CCACGGTTCAAGAAACTTGCTGACATGTATGGTGGAGGTGATGACATGgtgagcaagggc 3' | 5' ATTGTTTGTACTTGTCCAAAAACCAAGTTCACCCTGAAGTTCAacggtgcctgGGATCCg 3' |
| HindIII-mCherry | | 5' AAGCTTcttgtacagctcgtccatgcc 3' |
| Hind-loxp-EcoR | 5' AGCTTATAACTTCGTATAGCATACATTATACGAAGTTATG 3' | |
| NeoR | 5' AACTGCAGGACGAGGCAGC 3' | |
| polyA | 5' ATGCCTGCTCTTTACTGAAGG 3' | |
| CDH1-EXON3 | 5'GTTACAGGCATGAGACACTGT 3' | 5'ATAAGGAAGCTCAAGCATAGAC3' |

**Movie legends**

Movie 1. Timelapse movie of ESI017-CDH1:mCitrine-CDH2:mCherry-CAAX:mCerulean reporter cell line for 44.5 hours of MPS-LM treatment. Live imaging was performed using confocal microscopy. Cell membrane is shown in cyan (mCerulean-CAAX), CDH1 (E-cadherin) in green (mCitrine), and CDH2 (N-cadherin) in red (mCherry), and. Scale bar: 20 μm.
